# Transmission trees on a known pathogen phylogeny: enumeration and sampling

**DOI:** 10.1101/160812

**Authors:** Matthew Hall, Caroline Colijn

## Abstract

One approach to the reconstruction of infectious disease transmission trees from pathogen genomic data has been to use a phylogenetic tree, reconstructed from pathogen sequences, and annotate its internal nodes to provide a reconstruction of which host each lineage was in at each point in time. If only one pathogen lineage can be transmitted to a new host (i.e. the transmission bottleneck is complete), this corresponds to partitioning the nodes of the phylogeny into connected regions, each of which represents evolution in an individual host. These partitions define the possible transmission trees that are consistent with a given phylogenetic tree. However, the mathematical properties of the transmission trees given a phylogeny remain largely unexplored. Here, we describe a procedure to calculate the number of possible transmission trees for a given phylogeny, and we show how to uniformly sample from these transmission trees. The procedure is outlined for situations where one sample is available from each host and trees do not have branch lengths, and we also provide extensions for incomplete sampling, multiple sampling, and the application to time trees in a situation where limits on the period during which each host could have been infected are known. The sampling algorithm is available as an R package (STraTUS).

## Introduction

The use of genetic data to reconstruct a pathogen transmission tree (a graph representing who infected who in an epidemic) has been the subject of considerable interest in recent years. Many different approaches have been proposed, both phylogenetic and non-phylogenetic (Aldrin *et al*., 2011; Jombart *et al*., 2014; Skums *et al*., 2017). In phylogenetic approaches, a phylogenetic tree reconstructed from sequences for pathogens sampled in an epidemic will specify the order of the coalescences of lineages, and also, if its nodes are dated, the time at which these occurred. Some approaches further assume that internal nodes in the phylogeny correspond to transmission events (Lau *et al*., 2015; Mollentze *et al*., 2014; Morelli *et al*., 2012), which in a dated phylogeny specifies infection dates, while others do not (Didelot *et al*., 2014, 2017; Hall *et al*., 2015; Klinkenberg *et al*., 2017). In either case, a phylogeny on its own does not determine who infected who, and extra components are required to reconstruct transmission events.

The assumption of coinciding lineage coalescences and transmission events may be unwise, and in particular it does not take into account within-host pathogen diversity (Giardina *et al*., 2017; Ypma *et al*., 2013). Several approaches have been taken that do not make it, one of which is to note that if a phylogeny from a completely sampled outbreak has its nodes annotated with the hosts in which each lineage was present, the transmission tree is known (Didelot *et al*., 2014, 2017; Hall *et al*., 2015). In particular, Hall *et al*. (2015) noted that the set of transmission trees for a known phylogeny, with complete sampling and assuming transmission is a complete bottleneck, is equivalent to the set of partitions of its nodes with the property that each part of each partition contains at least one tip and the subgraph induced by the nodes in each part is connected. However, the mathematical properties of this space of partitions remain largely unexplored.

Here, we provide procedures for counting the total number of these partitions (and hence the total number of transmission trees) for a known phylogeny. We also give an algorithm that samples uniformly from the set of such partitions. Initially we assume that the phylogeny is binary, sampling is complete, that each host provided one sample, and that nothing is known about the timings of each infection, but we go on in the Appendix to individually relax each of these assumptions individually, and then relax them together.

The procedures outlined here may be useful to researchers wishing to explore the structure that the phylogeny imposes on transmission tree space, or alternatively to explore whether a candidate transmission event is firmly (mathematically) ruled out by a phylogeny or set of phylogenies. Uniform sampling from transmission trees on a phylogeny is rapid and could allow public health researchers who are reconstructing outbreaks a quick guide to some of the most frequently-occurring transmission events among all transmission trees consistent with a set of sequence data. We include some numerical applications of our sampling approach, comparing transmission trees on balanced vs unbalanced phylogenies, and comparing random uniformly sampled transmission trees to transmission trees inferred with the *TransPhylo* approach (Didelot *et al*., 2017).

## New Approaches

Here we describe how to count, and uniformly sample, transmission trees for a known phylogeny in the simplest case where the phylogeny is binary, each host in the transmission tree is sampled once and only once, and no time limits are placed on the duration of an infection.

Let the phylogeny 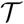 be an unlabelled rooted binary tree, without branch lengths. Let 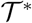 represent the unrooted tree obtained from 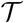 by attaching a single extra tip to the root of 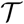 by a single edge. Note that two distinct 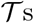 can have the same 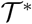, and that 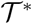 has one more tip than 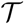.

We follow the correspondence described by Hall *et al*. (2015) between transmission trees and partitions of the node set of 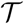 such that all tips derived from the same host are members of the same part (or block, or subset) of the partition, and the subgraph induced by each part is connected. This assumes that sampling is complete and that transmission is a complete bottleneck (i.e. that only one pathogen is transmitted at a time, so that diversity is not transmitted from host to host). While we relax the former assumption in the Appendix, the latter is more fundamental. See figure 1 for an example. We call a partition that satisfies these constraints an *admissible partition*.

**FIG. 1.**
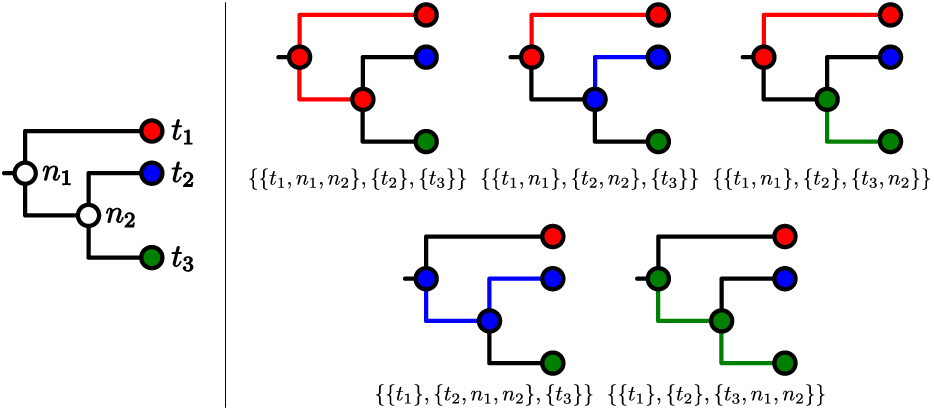
A rooted phylogeny (left) and the five compatible transmission trees labelled with their expression as partitions of its node set (right). Coloured branches connect members of the same part.

In this paper, the term “subtree” is intended in the normal phylogenetic (rather than graph theoretic) sense: a subtree is a subgraph of 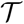 consisting of a node *u*, all its descendants (if any), and the edges between them. We denote the subtree rooted at *u* by 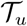; this is defined even if *u* is a tip.

Enumeration of possible transmission trees With 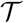 fixed and having *n* tips, suppose we wish to count the number of admissible partitions, as defined above, of its node set 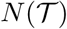, and hence the set of possible transmission trees. If the set of such partitions is 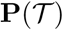, we wish to calculate 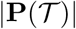. Nothing about the definition of an admissible partition requires a rooted tree, so 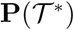 is defined similarly. It is trivial that if *n* = 1 then 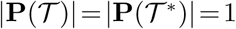. From here on, when we discuss partitions we mean admissible partitions.

If 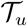 is a subtree, we can define 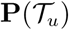 in the obvious way by regarding 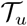 as a tree in its own right. If 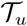 is indeed a subtree in a larger phylogeny of an epidemic, however, this is not sufficient. We do not assume that transmission occurs at the time of internal nodes, and so, even with complete sampling, it is possible that the root node of any subtree was not infecting any of the hosts from which the tips of that subtree were sampled.

To allow for this possibility, we also define a second set of partitions of 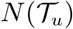, 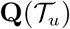:

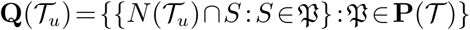

An element of 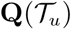 is the image of an element of 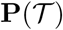 when the intersections of all its parts with the node set of 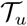 are taken. (This is not an injective operation, as the partition of the nodes of 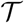 that are not nodes of 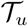 does not matter.)

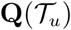, unlike 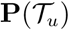, allows an internal node of 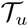 to share its part with no tip of 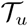. Suppose 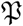 is a partition of 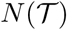 and there exists 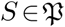 such that 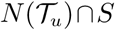 is nonempty and contains no tip of 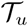. Then:

1. 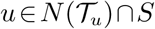 because if it were not then the *S* would not obey the connectedness requirement for being a part of a partition of 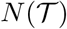. This is because, if 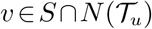 and *t* is the tip of 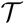 in *S*, then the path from *v* to *t* must intersect *u*.
2. 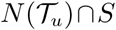 is the only member of the set 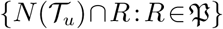 that contains no tips of 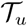, because *u* can belong to only one member of a partition of 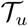.

It follows that 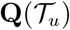 is the set of partitions of 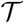 which obey the rules for an admissible partition except that they also allow (but do not insist on) an extra part (whose elements still induce a connected subgraph of 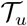) containing 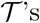 root. There is now no need to insist that 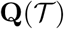 only be defined if 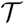 is a subtree of some larger tree; it is defined for any tree. Figure 2 shows an example of the extra elements of 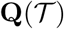 which are not already elements of 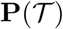 (and hence already displayed in figure 1).

**FIG. 2.**
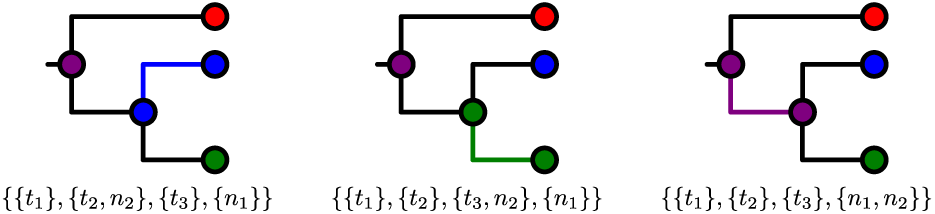
For the tree in figure 1, the three members of 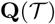 which are not members of 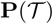.

We will not need to use the definition of 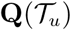 again, because it is in obvious correspondence with 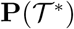. (Recall that 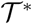 is obtained from 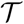 by attaching a single tip to 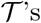 root.) Compare figure 3 with the full set of partitions displayed in figures 1 and 2 as an illustration of this.

**FIG. 3.**
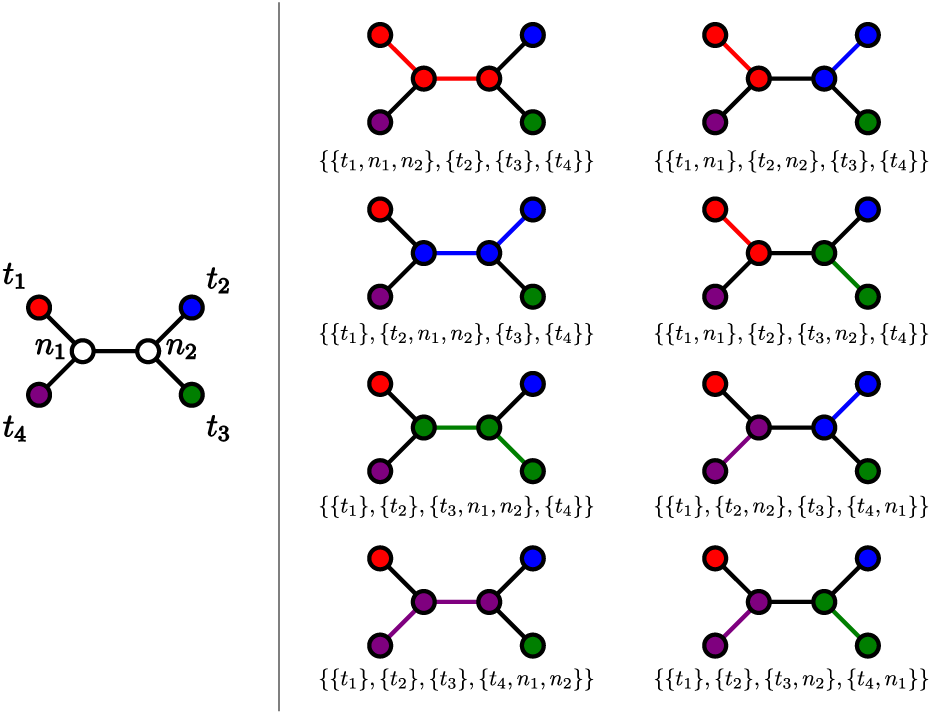
An unrooted phylogeny (left) and the eight partitions of its node set (right). Coloured branches connect members of the same part.

If *n* is at least 2, then 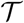 has a left subtree 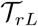 rooted at the left child *rL* of its root node r and a right subtree 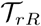 rooted at the right child *rR*.

### Proposition 1.

*If* 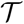 *has at least two tips, then:*

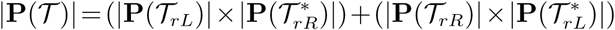

*Proof.*

Since 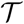 is not the tree with one node, its root *r* is internal. First we count the number of partitions where *r* is in the same part as a tip of 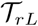. In this case, *rL* is in that same part by the connectedness requirement for parts: if it were not then the path from that tip to *r* would go through a node in a different part. The connectedness requirement then also insists that no node of 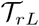 is in the same part as a tip of 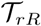, and the number of ways of partitioning the nodes of 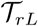 as a subtree of 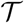 such that this is true is just 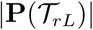. For each of those ways, the number of ways of partitioning the nodes of 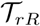 is 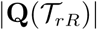, since some nodes of 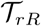 can be in the same part as *r* (and hence *rL*) and if any are then *rR* is. Since 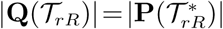 the number we are looking for is hence 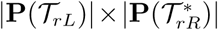.

An identical argument shows that the number of partitions where *r* is in the same part as a tip of 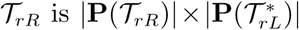, so the total number is 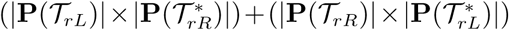.

### Proposition 2.

*If* 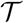 *has at least two tips, then:*

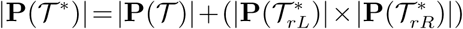

*Proof.*

The root *r* of 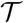 is an internal node of 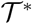 connected to a new tip, *t* (by the construction of 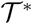). In some partitions of 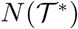, *t* is the only member of its part; there are obviously 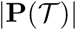 of these, because counting them is the same problem as counting partitions of the tree rooted at *r* with *t* excised.

If *t* is not the solo member of its part, *r* is in the same part by the connectedness requirement. The number of ways of partitioning 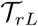 and 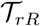 as subtrees of 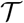 in this case are 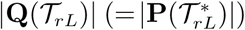 and 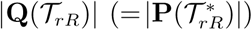 respectively. The total number of partitions for which *t* is not the only member of its part is the product of these.

Since 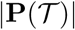 and 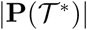 are equal to 1 when 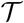 has one tip, 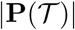 can now be calculated for any 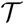 by doing a post-order tree traversal, as all that is needed to do the calculations at any node can be obtained by doing the same calculations at both of that node’s children. See figure 4 for an example.

### Proposition 3.

*If* 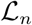 *is the ladder tree with n tips, then* 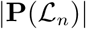 *is F*_2*n*–1_, *the* (2*n* − 1)*th Fibonacci number, and* 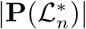 *is F*_2_*_n_*.

*Proof.*

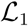 is the tree with one tip, so 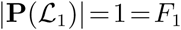 and 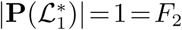, then proceed by induction. For *n*>1, the two subtrees descended from the root of 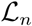 are 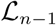 and 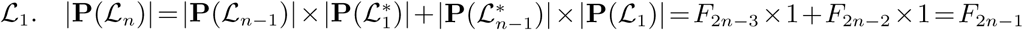 and 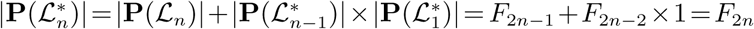.

### Enumeration of partitions with a known root part

Having demonstrated how to count the set of partitions or transmission trees compatible with a given 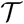, we now turn out attention to the matter of providing a uniform sample from that set. In order to do this, we need to determine what proportion of the 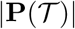 partitions have the root of 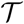 sharing its part with each tip.

If 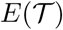 is the tip set of 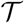, and 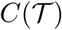 the set of children of *r*, let 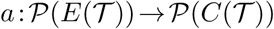 (with 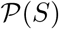 representing the power set of *S*) be the function taking a set of tips of 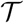 to the set of children of *r* which are ancestors of (or equal to) at least one of those tips.

**FIG. 4.**
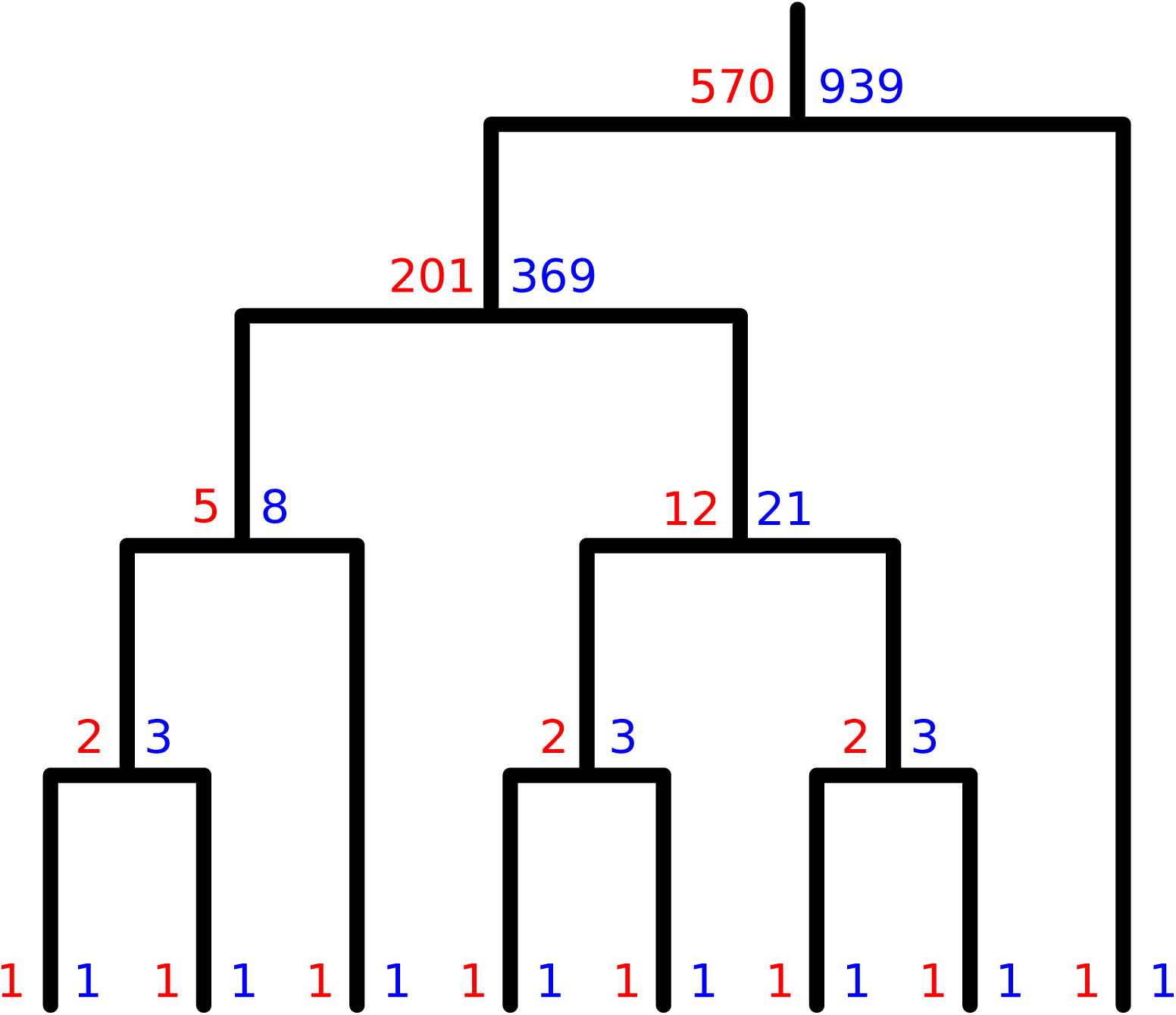
How to count partitions. At each node *u*, if 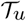 is the subtree rooted at *u*, then the red number is 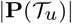 and the blue 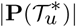. If *u* is internal and has children *uL* and *uR*, 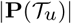 is 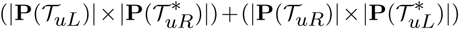 (the sum of the product of the blue number at *uL* and the red number at *uR*, and the product of the blue number at *uR* and the red number at *uL*), while 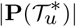 is 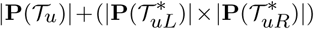 (the sum of the red number at *u* and the product of the blue numbers at its children).

Let {*t*_1_,…,*t_n_*} be the tips of 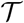. For each *i* let *H_i_* be the set containing just *t_i_*; this may seem redundant but it becomes crucial when relaxing the single sampling assumption as described in the Appendix. If 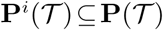 is the set of partitions of 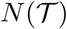 that have *r* in the same part as the membership of *H_i_*, we wish to calculate 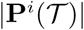 for all *i*. Naturally 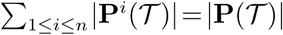. If 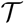 has one tip *t*_1_ ∈ *H*_1_, obviously 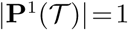. For any other 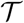, treating 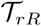 and 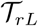 as trees in their own right but whose tips are partially shared with 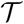, we can define 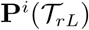 (respectively 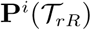) only if 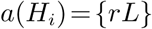 (respectively 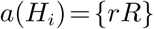).

#### Proposition 4.

*Suppose* 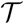 *has at least two tips*.

*Then:*

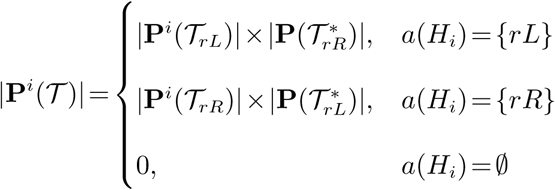

*Proof.*

Suppose *a*(*H_i_*) = {*rL*}. This forces *rL* to be in the same part as *r* by the connectedness requirement, and the number of members of 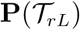 that have this property is 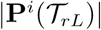. Such a choice of partition of 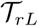 leaves 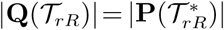 ways of partitioning the nodes of 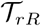. The argument if *a*(*t_i_*)=*rR* is identical. The final line is included for completeness in the case where 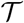 is a subtree of a larger tree and *t_i_* is a tip of the latter but not the former.

The value of 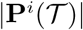 for all *i* can be calculated by a similar post-order traversal to that described in the previous section. See figure S1 for an example. Note that with an algorithm to calculate all 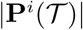, a separate one to calculate 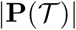 is not necessary as 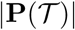 is just the sum of the 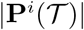.

### Sampling uniformly from 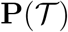

If the post-order traversal above is complete (and its results recorded for all subtrees of 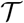, not merely 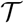 itself), sampling a random partition requires a single *pre*-order traversal. We start with a collection of empty sets 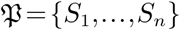, where each *S_i_* is to contain the set *H_i_*; once the traversal is complete, 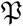 will be a partition of 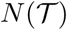. The traversal starts at *r*, and the 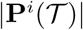 can be used as a set of probability weights for a draw of the *S_i_* that *r* belongs to, as they determine, for each *i*, how many of the 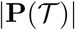 total partitions have *r* in *S_i_*.

Subsequently, when the traversal reaches another node *u* with parent *uP*, and we have already placed *uP* in *S_i_*, then *u* must also be placed in *S_i_* if *t_i_* is one of its descendants (by connectedness) or if *u* is *t_i_* itself. Otherwise, there are 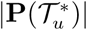 ways in which 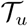 can be partitioned, since it can be a member of the same part as *uP* or a member of the same part as each of its tips. 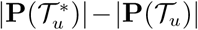 of these have *u* in the same part as *uP*, while the remaining 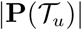 do not. For each *j* such that 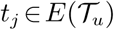, |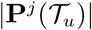 give the numbers of ways in which *u* can be placed in the same part as *t_j_*. The part for *u* can then be sampled with probability given by a weight vector that has 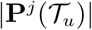 for each *S_j_* if 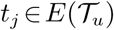, 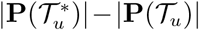 for *S_i_*, and 0 for any other part.

### Counting using the reduced Laplacian

Here we give an alternative non-recursive means of calculating 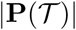 and 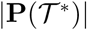, using graph theory. Following Levine (2009), let 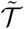 be the graph obtained from 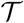 by collapsing all tips to a single node *s*.

We observe that there is a bijection between 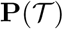 and the set of spanning trees of 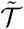. Suppose 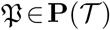 is a partition. The union of the subgraphs induced by each part of 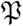 is a spanning forest of 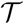 because every node belongs to one part. When 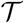 is transformed to 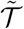, this spanning forest becomes a spanning tree: with the tips collapsed to a single node, the resulting graph is connected (because every part of 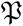 contains one tip) and has no cycles (because every part contains *only* one tip).

Conversely, if 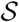 is a spanning tree of 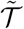 then, in “un-collapsing” the node *s* to recover 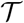, we split 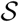 into *n* disconnected components. These clearly form a spanning forest which in turn defines a partition.

Let *u*_1,_…,*u_m_* be the nodes of 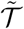, excluding *s* (although in fact, any single node can be excluded to give the same result), in an arbitrary order. If *d_i_* is the degree of *u_i_* and *d_ij_* the number of edges between *u_i_* and *u_j_* (universally 0 or 1 in our case), we now form the reduced Laplacian Δ of 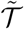, an *m* × *m* matrix:

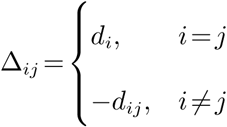

The matrix-tree theorem then states that the number of spanning trees of 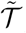, and hence the cardinality of 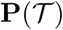, is the determinant of Δ.

The same procedure can be used to count the members of 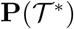, with 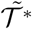 formed by collapsing the extra tip to *s* along with the rest. Levine (2009) defines 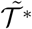 as the *wired tree* of 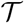, but the extra edge from 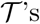 root to *s* specified in that definition is optional for our purposes.

As an example, if 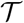 is the tree with three tips and *u*_1_ is its root, then the reduced Laplacian of 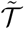 is 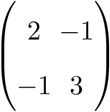, which has a determinant of 5, and that of 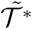 is 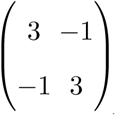 which has a determinant of 8, both as expected (see figures 1 and 3).

This result does not generalise to situations of incomplete or multiple sampling, where parts of a partition can include either zero or more than one tip: in the former case a spanning forest of 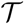, following the transformation to 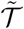, remains a forest rather than a tree, and in the latter case it has cycles. Thus, we leave further exploration of this avenue for future work.

## Results

Sampling random transmission trees that are consistent with a fixed phylogenetic tree has applications in transmission inference and in phylodynamics. In particular, there has been some work on whether imbalanced phylogenies are indicative of specific kinds of transmission (Colijn and Gardy, 2014; Frost and Volz, 2013; Leventhal *et al*., 2012; Robinson *et al*., 2013). It is clear that the phylogenetic tree places some constraints on who may have infected whom, particularly if individuals are treated and become uninfectious at the time of sampling. The current work aids investigations of this nature by permitting quantitative comparison of transmission trees sampled uniformly at random from two different phylogenetic trees.

The shapes of phylogeneties have been related to transmission patterns in a number of studies, as phylogenetic data are an appealing alternative to classical methods, such as contact tracing, to investigate transmission particularly in settings where highly-transmitting individuals may be difficult to identify directly, for example in sexually transmitted or blood-borne infections (Leventhal *et al*., 2012). In particular, how so-called “superspreaders” (individuals transmitting an infection to a large number of secondary cases), or contact number heterogeneity more broadly, may leave a signature in phylogenetic trees is one important phylodynamic application, particularly in HIV. Several studies have related contact number heterogeneity to the imbalance and cluster patterns in phylogenetic trees, with conclusions that differ depending on assumptions about the network structure and dynamics and the simulation approach (Colijn and Gardy, 2014; Frost and Volz, 2013; Leventhal *et al*., 2012; Robinson *et al*., 2013). One of the most commonly used ways to describe the shapes of phylogenetic trees is with their overall asymmetry (imbalance), via, for example, the Sackin index (Sackin, 1972). Indeed, in the phylodynamic literature, this and the number of cherries in the phylogeny have been the primary measures of tree shape. We explored whether there is a systematic difference in the offspring distribution in randomly sampled transmission trees resulting from their asymmetry.

We began with two input phylogenetic trees each with 40 tips, one from a Yule model (a pure branching process) and one from a so-called “biased” model with a bias parameter 0.9. The biased model is a growing tree model; the children of a lineage with a speciation rate *r* have rates *pr* and (1 − *p*)*r*. This produces imbalanced trees. The two input trees, along with a randomly sampled partition assuming full sampling and only one tip per host, are shown in Figure 5.

**FIG. 5.**
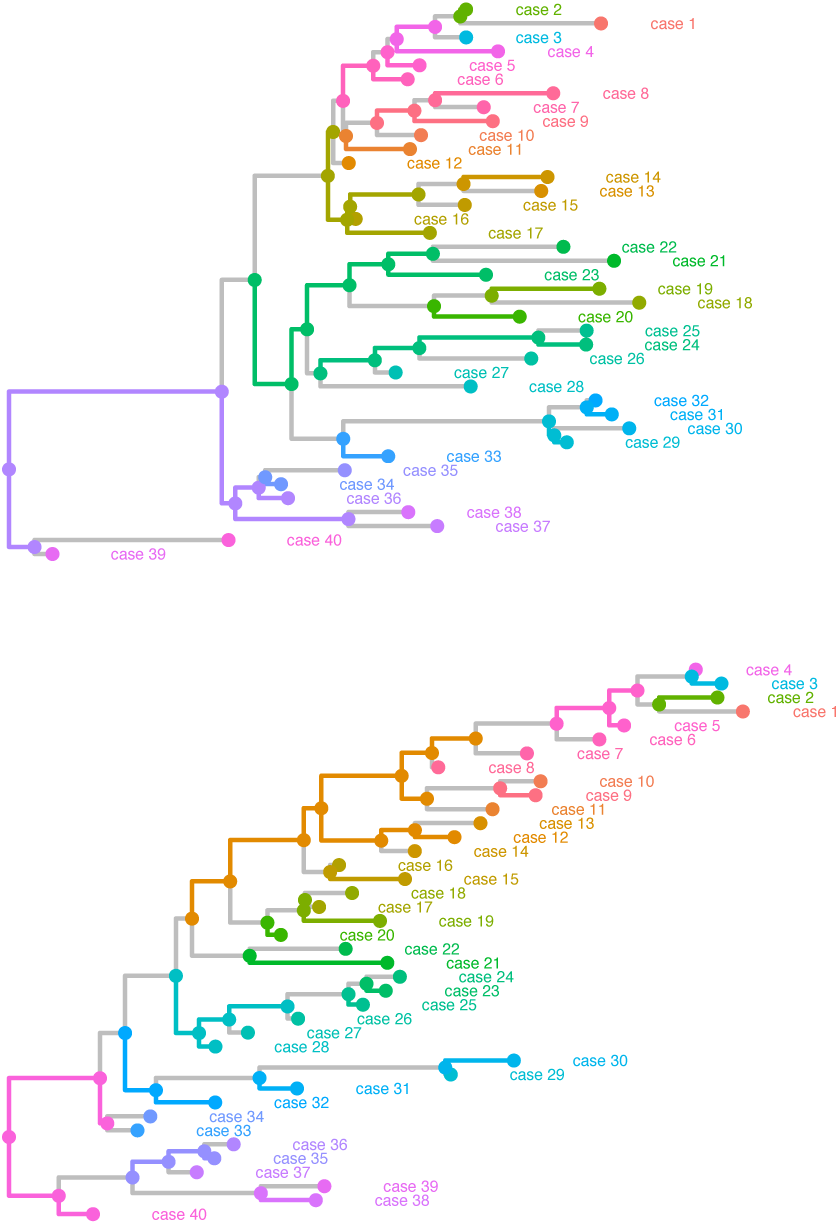
Yule (top) and biased (bottom) phylogenetic trees with randomly sampled partitions. Each colour corresponds to a part of each partition. Grey edges separate nodes that are in different parts of the partition.

We sampled 300 transmission trees uniformly at random on our two random trees, with full sampling and one tip per host, and compare the distribution of the number of secondary cases infected by a host (offspring distribution). With full sampling, the mean number of secondary cases in a tree is just under 1, because each individual except the source has a single infector.

We find that the relationship between the phylogenetic tree and the dispersion of the offspring distribution depends on whether the timings of infection are restricted. When we make no restrictions on the timing, there is sufficient flexibility in who may infect whom that the two trees have very similar offspring distributions. In contrast, if we constrain the heights of nodes in each tip’s part of the partition according to an infectious period (using the procedure outlined in the Appendix), the imbalanced tree is more constrained and has a less variable offspring distribution than the Yule tree (Figure 6).

**FIG. 6.**
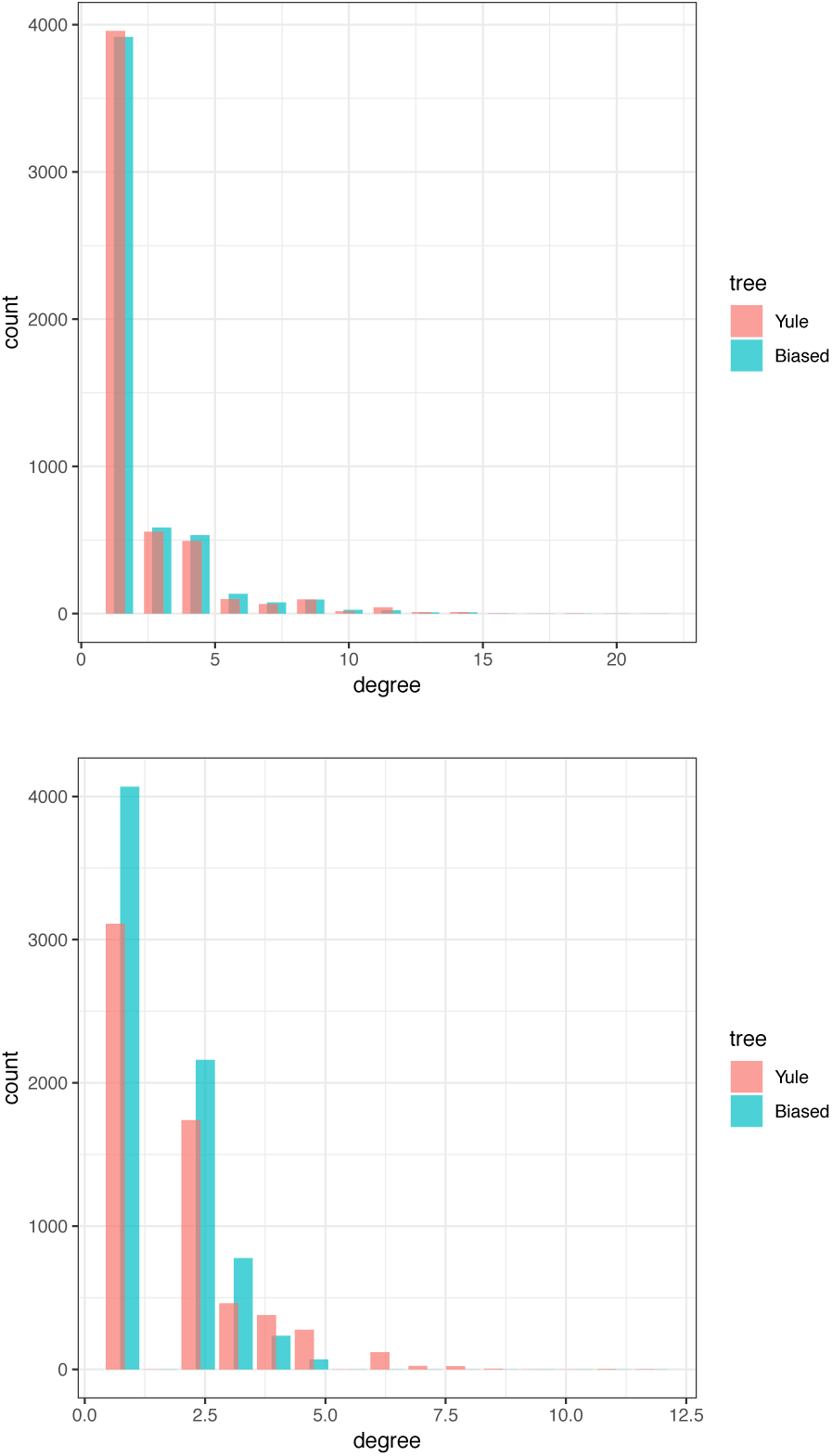
Offspring distributions from two input phyogenetic trees without (left) and with (right) constraints on the time between infection and sampling such that hosts became non-infectious immediately upon sampling, and had been infectious for a maximum of 3 (Yule) or 3.5 (biased) time units, compared to the mean branch length in these trees of 1.39 and 1.63 time units respectively.

We also compared the transmission trees sampled on the biased and Yule phylogenies directly, using the metric approach developed by Kendall *et al*. (2018). Briefly, the metric is a distance between two transmission trees; the distance is zero if and only if the transmission trees are the same (except for some sets of unsampled cases which are not relevant here, as we used full sampling). We compute the distances between all pairs of trees, and visualise the distances using principal component analysis (PCA). Figure 7 shows the results. The Yule and biased phylogenies both admit a “wide spread” of possible transmission trees, but they are strongly separated on the plot. This is a visual illustration of the fact that the structure of the phylogeny places consistent constraints on admissible transmission trees.

**FIG. 7.**
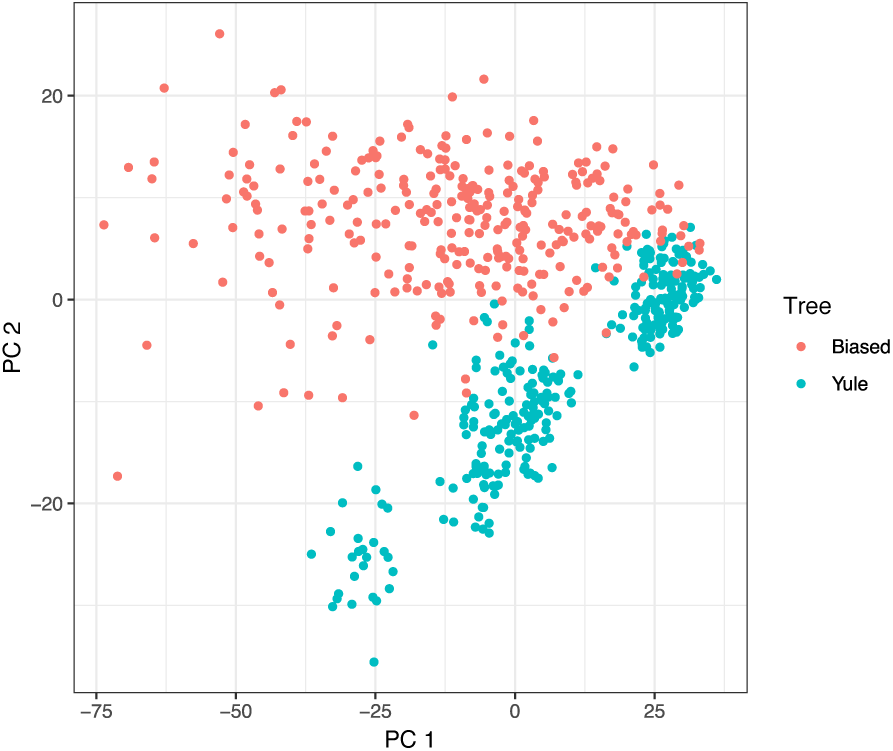
Principal component analysis plot visualising distances between transmission trees sampled on the Yule and biased phylogenies, without restrictions on the timing of transmission events.

We then sampled 500 random phylogenetic trees of 20 tips each and computed the number of transmission trees each one admits. We also computed two common tree shape statistics: the number of cherries and the Sackin imbalance. A cherry is a configuration consisting of two tips and an internal node. Each binary phylogenetic tree has at least one cherry and could have at most *n*/2 cherries. The Sackin imbalance (Blum and Francois, 2005; Sackin, 1972) has been defined in several ways, including the total or alternatively the average path length from a tip to the root of the tree. Broadly, the number of possible transmission trees on a fixed phylogeny increases as the Sackin imbalance increases, and declines as the number of cherries increases (cherries are symmetric feature, so trees with higher numbers of cherries tend to have a lower Sackin imbalance). This is, again, under the assumption that there no constraints on the timing of transmission relative to the node’s sampling time.

Finally, we compared randomly sampled transmission trees to transmission trees estimated by the *TransPhylo* algorithm (Didelot *et al*., 2017). We used an outbreak of tuberculosis cases over a 13 year period in Hamburg, Germany, which was previously published (Roetzer *et al*., 2013) and previously analysed in *TransPhylo* (Didelot *et al*., 2017). Because the current version of *STraTUS* cannot apply limits on infection times to unsampled cases (see the Appendix), we applied the two algorithms to a 72-tip subtree in which the root node of the epidemic was plausibly infecting a sampled host (see figure S2). (This restriction in *STraTUS* means it will not favour any particular number of unsampled hosts in the transmission tree separating the root node from the first sampled case, regardless of the branch lengths involved. This is very different to *TransPhylo*, so we ensure that the root case was plausibly sampled in order to make a comparison.) We sampled transmission trees uniformly at random with *STraTUS*, and compared them to the *TransPhylo-estimated* trees. The timed phylogeny was estimated in *BEAST* (Suchard *et al*., 2018) and was the same as reported in (Didelot *et al*., 2017), then pruned to the 72-tip subtree. We restricted the maximum possible time between infection and sampling to 7 years, and assumed that cases become noninfectious upon sampling. We generated multiple *STraTUS* samples for zero, 20, 40 and 60 unsampled hosts, and also with the unsampled count drawn from the empirical distribution of unsampled hosts from *TransPhylo*. The median number of unsampled hosts from *TransPhylo* was 39.

We used the metric and MDS approach outlined above to compare the sets of transmission trees. Figure 9 illustrates the results in 2-dimensional MDS. For 40 unsampled hosts and when the unsampled count was drawn from the *TransPhylo* empirical distribution, the *STraTUS* sample occupies much of the same space as *TransPhylo*, but that the *STraTUS* transmission trees are much more widely distributed. This is not surprising, as the sampling of a *TransPhylo* tree is determined by its posterior probability under a phylodynamic model, whereas *STraTUS* is a cruder, uniform sample from the space of all admissible phylogenies. The *STraTUS* sample with no unsampled hosts, on the other hand, forms a largely distinct cluster in the plot from the *TransPhylo* trees.

**FIG. 8.**
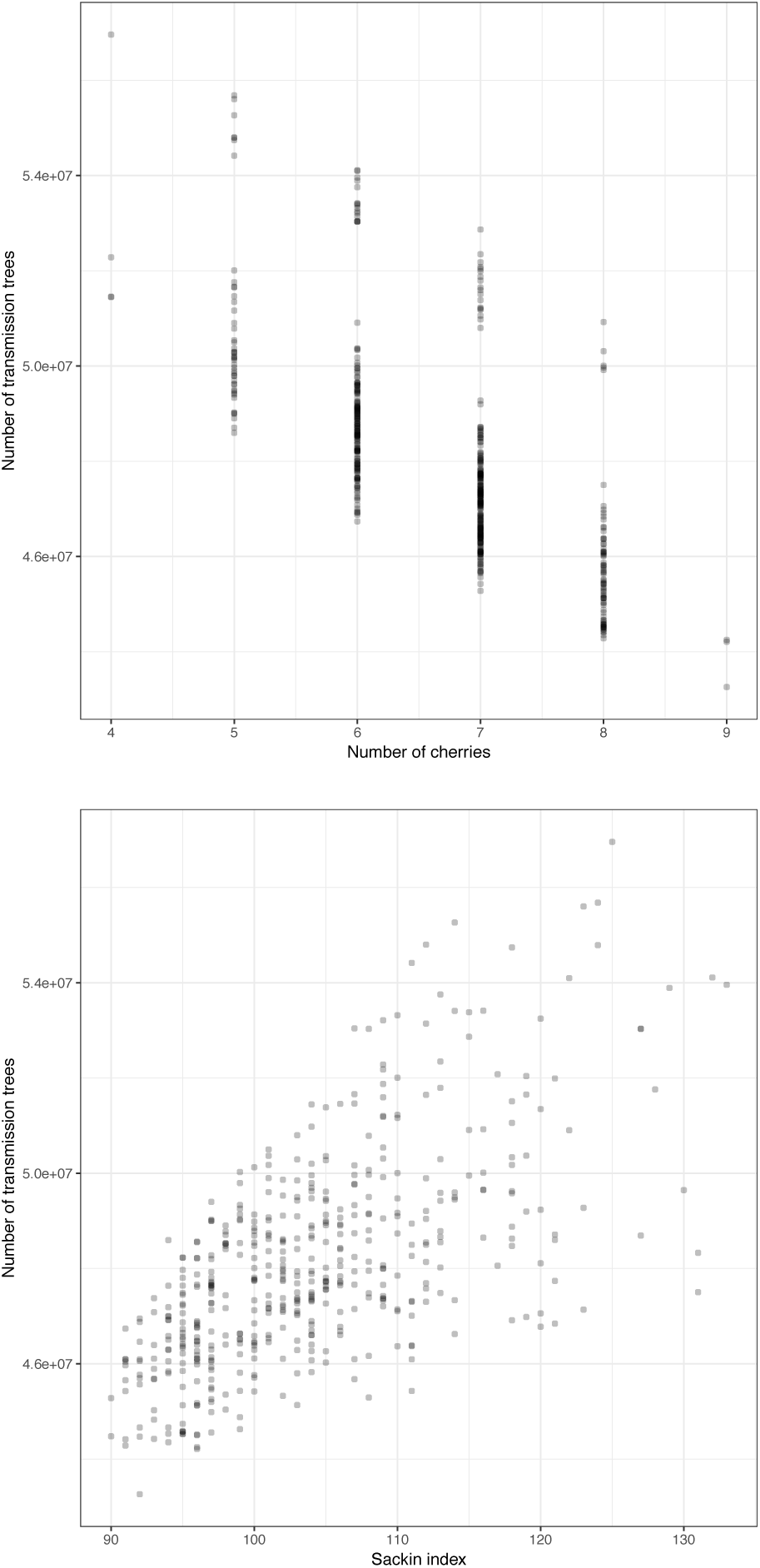
The number of transmission trees vs the number of cherries (left) and the Sackin measure of imbalance (right) over 500 random phylogenies each with 20 tips.

**FIG. 9.**
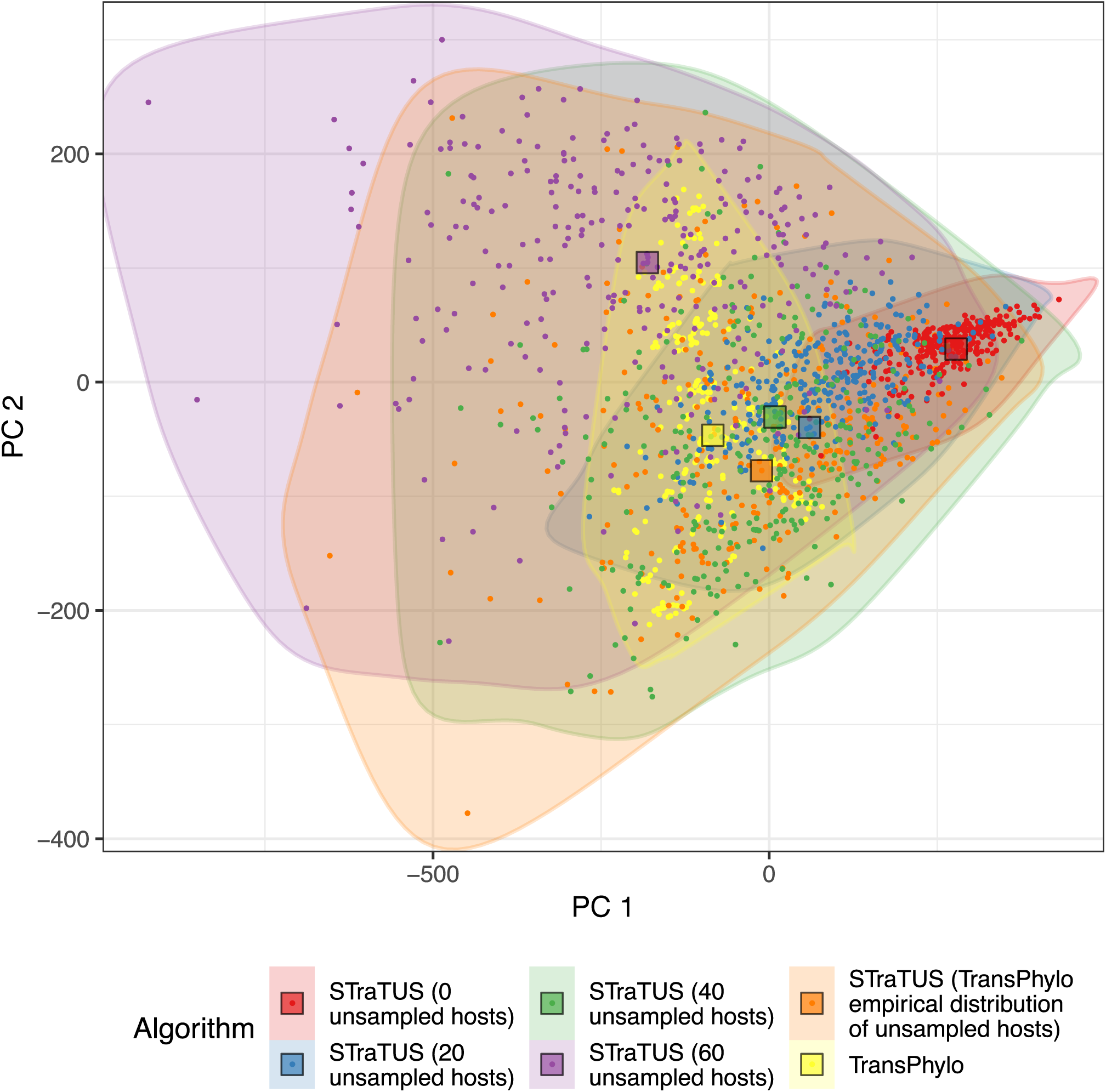
PCA plot illustrating the distances between transmission trees inferred by *TransPhylo* and sampled using *STraTUS*, derived from the timed phylogenetic tree of the Roetzer outbreak, previously used in (Didelot *et al*., 2017). The colours indicate the algorithm used and the number of unsampled cases selected in *STraTUS*. The shaded areas represent the extent of the MDS space occupied by the trees in each sample. The squares represent the geometric median tree of each sample.

We also determined the tree within each group that is closest to the centre of the trees (the geometric median tree; (Jombart *et al*., 2017)). These are marked on figure 9. Notably, the *STraTUS* sample whose median is closest to the *TransPhylo* median is the one where the unsampled host count was drawn from the *TransPhylo* empirical distribution. These results suggests that it may be possible to use *TransPhylo* to quickly produce an approximate sample of possible transmission trees for a given phylogeny, but that unbiased estimation of the number of unsampled individuals would be necessary.

For the *TransPhylo* and empirical *STraTUS* samples, we show the geometric median transmission trees in Figure 10. While these are different, they share a number of transmission events and features. The distribution of unsampled cases is notably different because *STraTUS* does not take branch lengths into account in placing them while *TransPhylo* does.

**FIG. 10.**
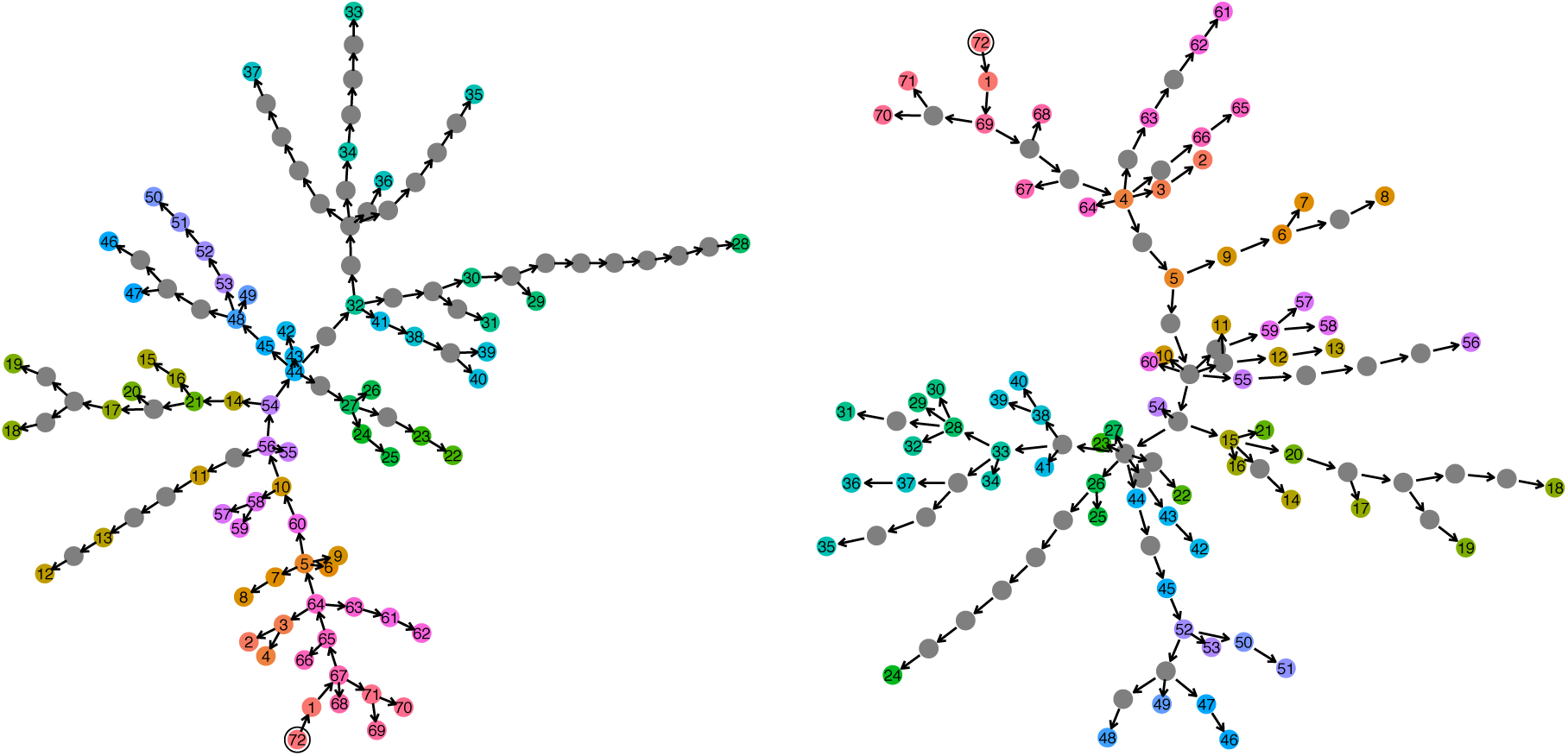
Geometric median trees from *TransPhylo* (left) and *STraTUS* (right). Grey nodes represent unsampled cases and in each case the index host in the tree is ringed in black. While many individual transmission events differ, there are many points where the differences are “minor” and the trees share small sub-clusters of cases who transmitted to each other in different configurations.

## Discussion

In this paper, we have explored the mathematics of the set of transmission trees for a known phylogeny, if internal nodes of that phylogeny are not taken to represent infection events, in greater depth and with more rigour than in any previous work. We also give algorithms for uniform sampling of transmission trees. In addition to establishing a firm footing for further theoretical work of this nature, and providing a new method to investigate the relationship between properties of the phylogeny and disease epidemiology, this work has several other potential applications.

The packages *TransPhylo* (Didelot *et al*., 2014, 2017) and *BEASTLIER* (Hall *et al*., 2015) both employ MCMC sampling of partitioned trees to estimate transmission trees, for a fixed phylogeny in the former case and a variable one in the latter. The uniform sampling procedure detailed here, perhaps together with recently-developed metrics on phylogeny and transmission tree space (Kendall and Colijn, 2016; Kendall *et al*., 2018) may prove valuable in the design of improved transition kernels for these algorithms. A uniform sampler for transmission trees may also be useful in a two-stage importance sampling approach of the type employed by Numminen *et al*. (2014), wherein a uniform sample of transmission trees are sampled, given importance weights according to their likelihood under a model of transmission and then resampled with probability proportional to those weights.

Furthermore, approaches such as *TransPhylo*, *BEASTLIER*, *phybreak* and others make use of a number of models and priors, including for example the natural history of the pathogen (which is used to inform the likelihood based on time between infection and transmission using a generation time), the sampling fraction and sampling process, and a coalescent model for the within-host pathogen evolution. These parameters are difficult to estimate in any single outbreak dataset (particularly in-host evolutionary parameters), and may vary from one outbreak or setting to the next. Re-using past estimates may not solve the problem. The ability to very rapidly sample from all transmission trees consistent with a phylogeny could allow outbreak investigators to rapidly understand which putative transmission events are and are not consistent with genomic data, without making strong assumptions on unknown parameters. When doing this, reasonably accurate quantification of the number of unsampled hosts would appear to be advisible.

The main assumption in transmission tree inference that we are unable to relax is the complete bottleneck at infection. The partition approach basically requires this, as to discard it is to discard the requirement that the region of a phylogeny associated with each host is connected. Without this, any number of transitions amongst the hosts can occur on any branch, and thus the set of transmission trees is infinite. We would argue that that set is actually rather less realistic than the one we present here, as large numbers of reinfection events will be rare for most pathogens. The importance of the bottleneck assumption in practice has not been extensively studied. Consequential violations of connectedness, where transmission trees exist that are actually impossible under the complete bottleneck assumption (see figure S3), require not just that multiple lineages be transmitted, but that two or more of them are later either transmitted onwards to different hosts, or sampled. How likely this is to happen in practice will vary from pathogen to pathogen and setting to setting; it is more plausible when the “hosts” in the transmission tree are premises rather than individual organisms. It is also unclear whether such an event would ever leave a sufficient signal on the pathogen genome to allow its identification. Nevertheless, methods to estimate transmission trees that do not make the complete bottleneck assumption are available (Worby *et al*., 2016).

## Materials and Methods

Random phylogenetic trees were generated using the `rtree` function in the `R` package *ape* v1.5 and all trees were visualised with *ggtree* v3.7. Transmission trees were compared using the metric of Kendall *et al*. (2018) implemented in *treespace* 1.1.3 (Jombart *et al*., 2017). Principal component analysis was performed using *ade4* v1.7-13 (Chessel *et al*., 2004). Sequencing, alignment and BEAST analysis of the *M. tubercolosis* dataset has been previously described (Didelot *et al*., 2017; Roetzer *et al*., 2013).

## Supporting information

## Acknowledgments

This work was supported by European Research Council (advanced grant number) PBDR-339251 and Engineering and Physical Science Research Council (UK) (grant numbers EP/K026003/1 and EPSRC EP/N014529/1).

