## Supplementary material for "Transmission trees on a known pathogen phylogeny: enumeration and sampling"

In this appendix, we extend the procedures for enumeration and sampling of transmission trees to multifurcating phylogenies, to situations of incomplete and/or multiple sampling, and to trees with node dates when restrictions on the timings of the infection of each host are specified.

### Multifurcating phylogenies

A modification to the enumeration and sampling algorithms is fairly trivial if $\mathcal{T}$ is not binary. If the root $r$ has $n$ children $rC_1, \ldots, rC_n$ then:

$$|\mathbf{P}(\mathcal{T})| = \sum_{1 \leq i \leq n} \Big( |\mathbf{P}(\mathcal{T}_{rC_i})| \prod_{\substack{1 \leq j \leq p \\ j \neq i}} |\mathbf{P}(\mathcal{T}^*_{rC_j})| \Big)$$

and

$$|\mathbf{P}(\mathcal{T}^*)| = |\mathbf{P}(\mathcal{T})| + \prod_{1 \leq i \leq n} |\mathbf{P}(\mathcal{T}^*_{rC_i})|$$

and the traversals can be performed in the same way as described in the main text.

### Incomplete sampling

If $\mathcal{T}$ is again binary, suppose that not every host in the transmission tree was sampled, but instead that there were $m$ unlabelled unsampled individuals, all of which are ancestral to at least one sampled individual. It is not sufficient merely add $m$ extra nonempty parts, each containing no tips, to a partition. This is because, to give a simple example, if an unsampled host $B$ was infected by a sampled host $A$ and directly infects only one other host $C$ which was also sampled, the region of the phylogeny corresponding to the infection of $B$ exists only along a branch (the branch whose parent node is in the part containing $A$'s tip and whose child node is in the one containing $C$'s tip) and no internal nodes are associated with $B$ at all. Nonetheless, the procedure for counting, and sampling from, the set of tree partitions with $n$

parts each containing a single tip and $m$ containing none turns out to lead to the more general answer as a byproduct of the calculations.

Suppose $\mathbf{P}_m(\mathcal{T})$ is the set of partitions of $\mathcal{T}$ with $n$ parts each containing a single tip and $m$ extra parts containing no tips. ($\mathbf{P}(\mathcal{T})$ as described above is $\mathbf{P}_0(\mathcal{T})$). Let $\mathbf{PS}_m(\mathcal{T})$ be the subset of $\mathbf{P}_m(\mathcal{T})$ where the root of $\mathcal{T}$ shares its part with a tip, and $\mathbf{PU}_m(\mathcal{T})$ the subset where it does not. $\mathbf{P}_m(\mathcal{T}^*)$ can be defined, although as $\mathcal{T}^*$ has no root $\mathbf{PS}_m(\mathcal{T}^*)$ and $\mathbf{PU}_m(\mathcal{T}^*)$ cannot be. $\mathbf{Q}_m(\mathcal{T})$ can also be defined and is exactly analogous to $\mathbf{P}_m(\mathcal{T}^*)$.

In a partition, call the parts containing tips the sampled parts, and those not containing tips the unsampled parts. If $\mathcal{T}$ has a single tip then $\mathbf{P}_m(\mathcal{T}) = \emptyset$ and $\mathbf{P}_m(\mathcal{T}^*) = \emptyset$ for all $m > 0$, because no internal nodes exist to be assigned to unsampled parts.

#### Enumeration of possible transmission trees

**Proposition 5.** *If $\mathcal{T}$ has at least two tips, then*

$$|\mathbf{PS}_m(\mathcal{T})| = \sum_{i=0}^{m} \left( (|\mathbf{P}_i(\mathcal{T}_{rL})| \times |\mathbf{P}_{m-i}(\mathcal{T}^*_{rR})|) + (|\mathbf{P}_{m-i}(\mathcal{T}_{rR})| \times |\mathbf{P}_i(\mathcal{T}^*_{rL})|) \right)$$

*Proof.* Since $r$ is not in an unsampled part, the $m$ such parts must be split between the subtree descended from $r$'s left child and that descended from its right child. The summation expresses the number of ways to make this split. Apart from this adjustment the argument is the same as in proposition 1.

□

**Proposition 6.** *If $\mathcal{T}$ has at least two tips, then*

$$|\mathbf{PU}_m(\mathcal{T})| = \sum_{i=0}^{m-1} (|\mathbf{P}_i(\mathcal{T}^*_{rL})| \times |\mathbf{P}_{m-1-i}(\mathcal{T}^*_{rR})|)$$

*Proof.* Since $r$ is in an unsampled part, one of the $m$ such parts is accounted for. The remaining $m-1$ are split amongst the left and right subtrees as above.

If we consider $\mathcal{T}_{rL}$ in isolation, we want to count the number of ways of partitioning its nodes with $i$ unsampled parts for certain and possibly one extra which, if it exists, must contain $rL$. (This would be the intersection of $N(\mathcal{T}_{rL})$ with a part $S$ of a member of $\mathbf{P}(\mathcal{T})$ such that $r \in S$.) This number is $|\mathbf{Q}_i(\mathcal{T}_{rL})| = |\mathbf{P}_i(\mathcal{T}^*_{rL})|$. Since exactly the same applies to $\mathcal{T}_{rR}$, the product of $|\mathbf{P}_i(\mathcal{T}^*_{rL})|$ and $|\mathbf{P}_{m-1-i}(\mathcal{T}^*_{rL})|$ is the desired number for a known $i$.

□

**Proposition 7.** *If $\mathcal{T}$ has at least two tips, then*

$$|\mathbf{P}_m(\mathcal{T}^*)| = |\mathbf{P}_m(\mathcal{T})| + \sum_{i=0}^{m} (|\mathbf{P}_i(\mathcal{T}^*_{rL})| \times |\mathbf{P}_{m-i}(\mathcal{T}^*_{rR})|)$$

*Proof.* Identical to proposition 2 except that we again allow for all the ways that the $m$ unsampled parts can be distributed.

□

Propositions 5 to 7 then allow us to calculate $|\mathbf{P}_m(\mathcal{T})| = |\mathbf{PU}_m(\mathcal{T})| + |\mathbf{PS}_m(\mathcal{T})|$ by a post-order traversal; note that at every internal node $u$ we must calculate $|\mathbf{P}_i(\mathcal{T}_u)|$ and $|\mathbf{P}_i(\mathcal{T}^*_u)|$ for all $i$ with $0 \leq i \leq m$, not just $|\mathbf{P}_m(\mathcal{T}_u)|$ and $|\mathbf{P}_m(\mathcal{T}^*_u)|$.

#### Enumeration of partitions with a known root part

Once again, let $t_1, \ldots, t_n$ be the tips of $\mathcal{T}$ and $H_1, \ldots, H_n$ a collection of sets containing just each $t_i$. Let $\mathbf{v}^i(\mathcal{T})$ be the vector of length $m+1$ whose $j$th entry is $|\mathbf{P}^i_j(\mathcal{T})|$, the number of partitions of $\mathcal{T}$ such that $r$ is in the same part as the members of $H_i$ if there are $j-1$ unsampled parts.

**Proposition 8.**

$$|\mathbf{P}^i_m(\mathcal{T})| = \begin{cases} \sum_{k=0}^{m} \left(|\mathbf{P}^i_k(\mathcal{T}_{rL})| \times |\mathbf{P}_{m-k}(\mathcal{T}^*_{rR})|\right) & a(H_i) = \{rL\} \\ \sum_{k=0}^{m} \left(|\mathbf{P}^i_k(\mathcal{T}_{rR})| \times |\mathbf{P}_{m-k}(\mathcal{T}^*_{rL})|\right) & a(H_i) = \{rR\} \\ 0 & a(H_i) = \emptyset \end{cases}$$

*Proof.* Analagous to proposition 4 after counting the ways the $i$ unsampled parts can be split between the two child subtrees. □

The values of $\mathbf{v}^i(\mathcal{T})$ for all tips $t_i$ of $\mathcal{T}$ can now be calculated by another post-order traversal.

#### Sampling uniformly from $\mathbf{P}_m(\mathcal{T})$

To start the pre-order traversal, we need a collection of sets $\mathfrak{P}$ which contains, initially, $n$ empty sets $\{S_1, \ldots, S_n\}$ which are each to contain a single tip, and $m$ other empty sets $\{U_1, \ldots, U_m\}$ which will not contain any tips. The procedure works by, at $r$, choosing a part using a vector of probability weights consisting of all the the $|\mathbf{P}^i_m(\mathcal{T})|$ for the sampled parts and $|\mathbf{PU}_m(\mathcal{T})|$ for the first unsampled part $U_1$. Once this is done, we must randomly choose how the remaining unsampled parts are divided between $\mathcal{T}_{rL}$ and $\mathcal{T}_{rR}$.

- If we chose $S_i$ for $r$ and $t_i$ is descended from $rL$, then the number of partitions which have $j$ unsampled parts amongst the nodes of $\mathcal{T}_{rL}$ (and hence $m-j$ amongst the nodes of $\mathcal{T}_{rR}$) is $|\mathbf{P}^i_j(\mathcal{T}_{rL})| \times |\mathbf{P}_{m-j}(\mathcal{T}^*_{rR})|$.
- If we chose $S_i$ for $r$ but instead $t_i$ is descended from $rR$, then the number of partitions which have $j$ unsampled parts amongst the nodes of $\mathcal{T}_{rL}$ is $|\mathbf{P}^i_{m-j}(\mathcal{T}_{rR})| \times |\mathbf{P}_j(\mathcal{T}^*_{rL})|$.
- If we chose $U_1$ for $r$, then the number of partitions which have $j$ *other* unsampled parts amongst the nodes of $\mathcal{T}_{rL}$ (and hence $m-1-j$ amongst the nodes of $\mathcal{T}_{rR}$) is $|\mathbf{P}_j(\mathcal{T}^*_{rL})| \times |\mathbf{P}_{m-1-j}(\mathcal{T}^*_{rR})|$.

In any case, we have a set of weights which we can use to randomly select $j$. This value is recorded for the number of previously unencountered unsampled parts in $\mathcal{T}_{rL}$; the remaining $m-j$ or $m-1-j$ are in $\mathcal{T}_{rR}$.

Now suppose the traversal has arrived at a node $u$ whose parent $uP$ has been assigned to a set $T$ (one of the $S_i$s or $U_i$s) and we know that there are $k$ previously unencountered unsampled parts in the partition of the nodes of $\mathcal{T}_u$. Then:

- If $T = S_i$ such that $t_i \in E(\mathcal{T}_u)$ or $t_i = u$ then we must place $u$ in $T$.
- Otherwise, we randomly choose a part for $u$ using the following weights:
  - $|\mathbf{P}^j_k(\mathcal{T}_u)|$ for $S_j$ if $t_j \in E(\mathcal{T}_u)$.
  - $|\mathbf{P}_k(\mathcal{T}^*_u)| - |\mathbf{P}_k(\mathcal{T}_u)|$ for the part that contains $uP$ (whether that is sampled or unsampled).
  - $|\mathbf{PU}_k(\mathcal{T}_u)|$ for $U_l$ where $l$ is the smallest index such that $U_l$ is still empty.

– 0 for any other part.

Then, if $u$ is not a tip, we split the $k-1$ (if we assigned $u$ to a new unsampled part) or $k$ (if we did not) remaining unencountered unsampled parts between $u$'s left and right descendant subtrees, as above with a slight modification:

- If we chose $S_i$ for $u$ and $t_i$ is descended from $rL$, then the number of partitions which have $j$ unsampled parts amongst the nodes of $\mathcal{T}_{rL}$ (and hence $k-j$ amongst the nodes of $\mathcal{T}_{rR}$) is $|\mathbf{P}^i_j(\mathcal{T}_{rL})| \times |\mathbf{P}_{k-j}(\mathcal{T}^*_{rR})|$.
- If we chose $S_i$ for $u$ and $t_i$ is descended from $rR$, then that number is $|\mathbf{P}^i_{k-j}(\mathcal{T}_{rR})| \times |\mathbf{P}_j(\mathcal{T}^*_{rL})|$.
- If we chose $S_i$ for $u$ and $t_i$ is descended from neither $rR$ nor $rL$, then that number is $|\mathbf{P}_j(\mathcal{T}^*_{uL})| \times |\mathbf{P}_{k-j}(\mathcal{T}^*_{uR})|$.
- If we chose $U_i$ for $u$ and $uP \in U_i$ (so we did not add $u$ to a previously empty set), then that number is also $|\mathbf{P}_j(\mathcal{T}^*_{uL})| \times |\mathbf{P}_{k-j}(\mathcal{T}^*_{uR})|$.
- If we chose $U_i$ for $u$ and $uP \notin U_i$, then that number is $|\mathbf{P}_j(\mathcal{T}^*_{rL})| \times |\mathbf{P}_{k-1-j}(\mathcal{T}^*_{rR})|$ as one of the $k$ unsampled elements is now accounted for.

##### Sampling uniformly from the set of transmission trees with $m$ unsampled hosts

To complete the picture, we now must consider the case where only $l$ of the $m$ unsampled hosts actually correspond to parts, and the remaining $m-l$ appear only along branches. If we have sampled an element $\mathfrak{P}$ of $\mathbf{P}_l(\mathcal{T})$ as above, we need a distribution of the additional $m-l$ hosts. Because we assume no superinfection, these hosts must appear on a branch whose endpoints are in different parts, or alternatively above the root node if they have no sampled ancestors in the transmission tree. There are $l+n$ such positions in the tree - the roots of the subgraphs induced by each part of $\mathfrak{P}$. Thus we are assigning $m-l$ identical objects to $l+n$ possibly empty groups, and the number of ways of doing this is $\binom{m+n-1}{l+n-1}$.

If we calculate $|\mathbf{P}_l(\mathcal{T})|$ for $0 \leq l \leq m$, then we know that there are $|\mathbf{P}_l(\mathcal{T})| \times \binom{m+n-1}{l+n-1}$ transmission trees with $m$ unsampled hosts where $l$ of those hosts have parts. We can randomly select an $l$ using those counts as probability weights, randomly generate an element of $\mathbf{P}_l(\mathcal{T})$ as above, and finally randomly assign the extra $m-l$ hosts to branches.

### Multiple sampling

Removing unsampled hosts from consideration, we now relax the assumption that each part of a partition contains only a single tip. Fix a partition $\mathfrak{H}$ of the tip set $E(\mathcal{T})$ of $\mathcal{T}$; this, in contrast to the partitions discussed thus far, and only divides the set of *tips* according to which host each corresponds to. We now investigate the set $\mathbf{P}(\mathcal{T};\mathfrak{H})$ which is the set of partitions $\mathfrak{P}$ of $N(\mathcal{T})$ such that $\{A \cap E(\mathcal{T}) : A \in \mathfrak{P}\} = \mathfrak{H}$ (i.e. the partitions of $N(\mathcal{T})$ which agree with $\mathfrak{H}$ on the tips). These correspond to the possible transmission trees, under the assumption that each host was only infected once (but since hosts may have provided more than one sample to the study, they may have more than one tip in the tree). Each part of $\mathfrak{H}$ contains all the tips sampled from a single host. $\mathbf{P}(\mathcal{T})$ as in the main text is $\mathbf{P}(\mathcal{T};\mathfrak{I})$ where $\mathfrak{I}$ is the partition of singletons of $E(\mathcal{T})$.

For a set $H \in \mathfrak{H}$ (i.e. a subset of tips corresponding to pathogens isolated from a single host) define the bridge $\mathbf{b}(H)$ of $H$ to be the minimal subset of $N(\mathcal{T})$ such that $H \subseteq \mathbf{b}(P)$ and the subgraph of $\mathcal{T}$ induced by $\mathbf{b}(P)$ is connected. This contains all elements of $H$, the MRCA of $H$, and all nodes on the paths between them. Obviously if $|H| = 1$ then $\mathbf{b}(H) = H$. See figure S4 for an example.

If any two elements of $\mathfrak{H}$ have bridges whose intersections are nonempty, then $|\mathbf{P}(\mathcal{T}; \mathfrak{H})| = 0$; there are simply no possible transmission trees because the connectedness requirement would insist that some nodes be part of more than one partition element. So assume that this is not true. For any partition of $N(\mathcal{T})$ which obeys the connectedness rules, being a member of a bridge forces a node to be a member of the same part as those tips whose bridge it belongs to. Define all the nodes that are members of any bridge as bridge nodes.

The intuitive way to consider $\mathbf{P}(\mathcal{T}_u; \mathfrak{H})$ for a subtree $\mathcal{T}_u$ of $\mathcal{T}$ rooted at $u$ would be to use the set $\mathfrak{H}_u = \{A \cap E(\mathcal{T}_u) : A \in \mathfrak{H}\}$ as a partition of the tips of $\mathcal{T}_u$. Bridge nodes determined by $\mathfrak{H}$ would not necessarily be determined as such by $\mathfrak{H}_u$. Unfortunately, this is not useful for our purposes, and we must retain the restrictions on the parts to which a node of $\mathcal{T}_u$ can belong that are determined by $\mathfrak{H}$, even when we move to counting partitions of $\mathcal{T}_u$. For example, if $H \in \mathfrak{H}$ consists of two tips $t_1$ and $t_2$, but only $t_1$ is a tip of $\mathcal{T}_u$, then in enumerating and sampling partitions we still consider the intersection $\mathbf{b}(A) \cap N(\mathcal{T}_u)$ to be bridge nodes, even though $\{t_1\}$ is a singleton element of $\mathfrak{H}_u$. (In fact $u$ must be a member of $\mathbf{b}(A)$.) So for any node $u$ of $\mathcal{T}$, we define $\mathbf{P}(\mathcal{T}_u; \mathfrak{H})$ to be the number of ways to partitioning $\mathcal{T}$'s nodes that respects the restrictions that $\mathfrak{H}$ places on which internal nodes can share elements with which tips. In summary:

- $\mathbf{P}(\mathcal{T}; \mathfrak{H})$ is the set of partitions of $N(\mathcal{T})$ such that if $\mathfrak{P} \in \mathbf{P}(\mathcal{T}; \mathfrak{H})$, each $S \in \mathfrak{P}$ contains one and only one part of $\mathfrak{H}$ and the subgraph induced by $S$ is connected. The connectedness requirement insists that if $S$ contains $P \in \mathfrak{H}$ then it in fact contains $\mathbf{b}(P)$.

- If $u$ is a node of $\mathcal{T}$ and $\mathcal{T}_u$ the subgraph rooted at $u$, $\mathbf{P}(\mathcal{T}_u; \mathfrak{H})$ is the set of partitions of $N(\mathcal{T}_u)$ such that if $\mathfrak{P} \in \mathbf{P}(\mathcal{T}_u; \mathfrak{H})$, each $S \in \mathfrak{P}$ contains $\mathbf{b}(P) \cap N(\mathcal{T}_u)$ for one and only one part $H$ of $\mathfrak{H}$ and the subgraph induced by $S$ is connected.

It should be fairly obvious that if an internal node is a bridge node then one or both of its children must be as well. $\mathbf{P}(\mathcal{T}^*; \mathfrak{H})$ has the definition one would expect, with the extra node forming a singleton extra part of $\mathfrak{H}$. For a node $u$, $\mathbf{P}(\mathcal{T}^*_u; \mathfrak{H})$ again respects the bridge nodes imposed by $\mathfrak{H}$ on $\mathcal{T}$.

Now suppose $\mathcal{T}$ has at least two tips and let $u$ be any internal node of $\mathcal{T}$, including $r$. Its children are $uL$ and $uR$.

**Proposition 9.**

$$|\mathbf{P}(\mathcal{T}^*_u; \mathfrak{H})| = \begin{cases} |\mathbf{P}(\mathcal{T}_u; \mathfrak{H})| + (|\mathbf{P}(\mathcal{T}^*_{uL}; \mathfrak{H})| \times |\mathbf{P}(\mathcal{T}^*_{uR}; \mathfrak{H})|), & u \textit{ is a not a bridge node} \\ |\mathbf{P}(\mathcal{T}_u; \mathfrak{H})|, & u \textit{ is a bridge node} \end{cases}$$

*Proof.* If $u$ is a bridge node then it cannot belong to the part containing the extra node, so the number of partitions is exactly the number with that node excised. Otherwise, the argument is as proposition 2. □

**Proposition 10.**

$$|\mathbf{P}(\mathcal{T}_u; \mathfrak{H})| = \begin{cases} (|\mathbf{P}(\mathcal{T}_{uL}; \mathfrak{H})| \times |\mathbf{P}(\mathcal{T}^*_{uR}; \mathfrak{H})|) + (|\mathbf{P}(\mathcal{T}_{uR}; \mathfrak{H})| \times |\mathbf{P}(\mathcal{T}^*_{uL}; \mathfrak{H})|), & u \textit{ is not a bridge node} \\ |\mathbf{P}(\mathcal{T}^*_{uL}; \mathfrak{H})| \times |\mathbf{P}(\mathcal{T}^*_{uR}; \mathfrak{H})|, & u \textit{ is a bridge node} \end{cases}$$

*Proof.* If $u$ is a bridge node then at least one of its children is too. At least one of those children, in fact, must be part of the same bridge as itself; suppose this is $uL$. Now $u$ and $uL$ must be in the same part so the $|\mathbf{P}(\mathcal{T}_{uL}); \mathfrak{H}|$ partitions of $uL$ determine which part $u$ belongs to, and because $uL$ is a bridge node $|\mathbf{P}(\mathcal{T}_{uL}); \mathfrak{H}| = |\mathbf{P}(\mathcal{T}^*_{uL}); \mathfrak{H}|$ by proposition 9. If $uR$ is not a bridge node then there are $|\mathbf{P}(\mathcal{T}^*_{uR};\mathfrak{H})|$ ways of partitioning the nodes of $\mathcal{T}_{uR}$ by a by now very familiar logic. If it is, then there are $|\mathbf{P}(\mathcal{T}_{uR},\mathfrak{H})|$ partitions of those nodes since the part that $uR$ belongs to is fixed, but $|\mathbf{P}(\mathcal{T}_{uR};\mathfrak{H})| = |\mathbf{P}(\mathcal{T}^*_{uR};\mathfrak{H})|$ by proposition 9. Obviously the same applies with $uR$ and $uL$ reversed. If $u$ is not a bridge node then the argument of proposition 1 still applies. □

If $\mathfrak{H}$ has $l$ parts, number them $H_1,\ldots,H_l$. For a subgraph $\mathcal{T}_u$, let $\mathbf{P}^i(\mathcal{T}_u,\mathfrak{H}) \subseteq \mathbf{P}(\mathcal{T}_u,\mathfrak{H})$ be those partitions where $u$ shares a part with the intersection $E(\mathcal{T}_u)\cap H_i$; this subset may be empty when that intersection itself is empty. Note that this definition works if $u = r$. As before, let the function $a$ take a set of tips to the set of children of $u$ that are ancestors of those tips.

**Proposition 11.**

$$|\mathbf{P}^i(\mathcal{T}_u,\mathfrak{H})| = \begin{cases} |\mathbf{P}^i(\mathcal{T}_{uL};\mathfrak{H})| \times |\mathbf{P}(\mathcal{T}^*_{uR};\mathfrak{H})|, & a(H_i)=\{rL\} \text{ and } u \notin \mathbf{b}(H_j) \text{ for } j \neq i \\ |\mathbf{P}^i(\mathcal{T}_{uR};\mathfrak{H})| \times |\mathbf{P}(\mathcal{T}^*_{uL};\mathfrak{H})|, & a(H_i)=\{rR\} \text{ and } u \notin \mathbf{b}(H_j) \text{ for } j \neq i \\ |\mathbf{P}^i(\mathcal{T}_{uL};\mathfrak{H})| \times |\mathbf{P}^i(\mathcal{T}_{uR};\mathfrak{H})|, & a(H_i)=\{rL,rR\} \\ 0, & u \in \mathbf{b}(H_j) \text{ with } j \neq i \end{cases}$$

*Proof.* First note that if $a(H_i) = \{rL, rR\}$ then $u$ is a bridge node and in particular a member of $\mathbf{b}(H_i)$. If $u$ is not a bridge node at all, we must be in one of the first two cases. The argument in that case follows that of proposition 4.

If $u \in \mathbf{b}(H_i)$ but only one child of $u$ has any descendant tips that are members of $H_i$, then suppose $uL$ is the child that does. Then $uR$ is either not a bridge node or a member of $\mathbf{b}(H_j)$ for $j \neq i$. The number of ways of partitioning the nodes of $\mathcal{T}_{uL}$ with $uL$ sharing a part with the members of $H_i$ is $|\mathbf{P}^i(\mathcal{T}_{uL},\mathfrak{H})|$ and, because $u$ shares a part with $uL$, each of those means $|\mathbf{P}(\mathcal{T}^*_{uR};\mathfrak{H})|$ ways of partitioning the nodes of $\mathcal{T}_{uR}$. The same goes with $uL$ and $uR$ reversed.

If both $uL$ and $uR$ have descendant tips that are members of $H_i$ then the number of partitions with $u$ sharing a part with the members of $H_i$ is just the product of the number of ways of partitioning both child subtrees in the same way. The nodes $u$, $uL$ and $uR$ must all be in the same part.

In any other situation $u \in \mathbf{b}(P_j)$ for $j \neq i$ and there cannot be any partitions that have it sharing a part with the members of $H_i$.

□

If $t$ is a tip then $|\mathbf{P}(\mathcal{T}^*_t;\mathfrak{H})| = 1$, $|\mathbf{P}(\mathcal{T}_t;\mathfrak{H})| = 1$, and $|\mathbf{P}^i(\mathcal{T}_t,\mathfrak{H})| = 1$ if $t \in H_i$ and 0 otherwise. This is all that is necessary to set up traversals analogous to those described under New Approaches.

### Infection time limits

Returning once more to the case where sampling is single and complete, we now give $\mathcal{T}$ branch lengths, which means a height function $h : N(\mathcal{T}) \to \mathbb{R}^+$ can be defined such that for all nodes $u$ with parent $uP$, $h(u) < h(uP)$. Branch lengths and heights are intended to be in units of calendar time, not genetic distance. We extend $h$ to $N(\mathcal{T}^*)$ by setting $h(t) = \infty$ if $t$ is the extra tip; while $\mathcal{T}^*$ remains formally unrooted, there is only one way to display it that makes sense.

Each set $H_i$, which for now once more contains just the tip $t_i$, is now associated with a closed interval $I_i = [\alpha_i, \beta_i]$ such that, for any partition $\mathfrak{P}$ of the nodes of $\mathcal{T}$, if $\{u\} \cup H_i \subseteq S_i \in \mathfrak{P}$ then $h(u) \in I_i$. (Obviously no partitions exist without $h(t_i) \in I_i$ for all $i$.)

The $I_i$s determine minimum and maximum heights for all the nodes in each part. This is useful if infection is expected to end with sampling, or if a maximum time from infection to sampling is known. If $\mathbf{I}$ is the complete set of intervals, then let $\mathbf{P}(\mathcal{T};\mathbf{I})$ by the set of partitions subject to these additional restrictions. For a subtree $\mathcal{T}_u$, $\mathbf{P}(\mathcal{T}_u;\mathbf{I})$ is the set of partitions subject to the restrictions where they are appropriate (i.e. the $I_i$ where $H_i$ actually contains tips of $\mathcal{T}_u$).

Let $\mathbf{P}(\mathcal{T}^*;\mathbf{I}\cup[\gamma,\infty))$ be the set of partitions of $\mathcal{T}^*$ such that the part containing the extra tip contains only nodes of heights greater than $\gamma$, and the restrictions imposed by $\mathbf{I}$ still apply to the other parts.

**Proposition 12.** *If $\mathcal{T}$ has at least two tips, then*

$$|\mathbf{P}(\mathcal{T}^*;\mathbf{I}\cup[\gamma,\infty))| = \begin{cases} |\mathbf{P}(\mathcal{T};\mathbf{I})|, & h(r)<\gamma \\ |\mathbf{P}(\mathcal{T};\mathbf{I})| + (|\mathbf{P}(\mathcal{T}^*_{rL};\mathbf{I}\cup[\gamma,\infty))| \times |\mathbf{P}(\mathcal{T}^*_{rR};\mathbf{I}\cup[\gamma,\infty))|), & h(r)\geq\gamma \end{cases}$$

*Proof.* If $h(r)<\gamma$ then $r$ cannot be in the same part as the extra tip, so the number of partitions is the same as in the rooted case. Otherwise, see proposition 2. □

A particularly concise expression for $|\mathbf{P}(\mathcal{T};\mathbf{I})|$ does not exist, so instead we suggest calculating it by first determining the number of partitions that have $r$ in the same part as each tip and then adding those up. So let $\mathbf{P}^i(\mathcal{T},\mathbf{I})$ be the set of partitions of $\mathcal{T}$ where $r$ is in the same part as the members of $H_i$ and the restrictions imposed by $\mathbf{I}$ apply.

**Proposition 13.** *If $\mathcal{T}$ has at least two tips, then*

$$|\mathbf{P}^i(\mathcal{T},\mathbf{I})| = \begin{cases} 0 & h(r)\notin I_i \\ |\mathbf{P}^i(\mathcal{T}_{rL},\mathbf{I})| \times |\mathbf{P}(\mathcal{T}^*_{rR};\mathbf{I}\cup[\alpha_i,\infty))|, & h(r)\in I_i \text{ and } a(H_i)=\{rL\} \\ |\mathbf{P}^i(\mathcal{T}_{rR},\mathbf{I})| \times |\mathbf{P}(\mathcal{T}^*_{rL};\mathbf{I}\cup[\alpha_i,\infty))|, & h(r)\in I_i \text{ and } a(H_i)=\{rR\} \end{cases}$$

*Proof.* If $h(r)$ lies outside $I_i$ then the answer is trivially zero. If not, and $t_i$ is descended from $rL$, then there are $|\mathbf{P}^i(\mathcal{T}_{rL};\mathbf{I})|$ ways of partitioning the nodes of $\mathcal{T}_{rL}$ such that $rL$ is in the same part as $t_i$. For each of these, we need the the number of ways of partitioning the nodes of $\mathcal{T}_{rR}$ such that an extra part may include the root, but that that part cannot contain any nodes whose heights are smaller than the lower limit of $I_i$, i.e. $\alpha_i$. This is $|\mathbf{P}(\mathcal{T}^*_{rR};\mathbf{I}\cup[\alpha_i,\infty))|$. As usual, an identical argument applies with $rL$ and $rR$ reversed. □

Then $|\mathbf{P}(\mathcal{T};\mathbf{I})| = \sum_{i=1}^{n} |\mathbf{P}^i(\mathcal{T};\mathbf{I})|$. If $\mathcal{T}$ has one tip, then both $\mathbf{P}(\mathcal{T};\mathbf{I})$ and $\mathbf{P}(\mathcal{T}^*;\mathbf{I}\cup[\gamma,\infty))$ (regardless of $\gamma$) have size 1 if the tip lies within its own interval and 0 if it does not. As in the previous section, the traversals described previously can be used to count the full set of partitions and sample uniformly from it.

### A unified counting and sampling procedure

We now relax all the assumptions at once. $\mathcal{T}$ may not be binary, there are $m$ unsampled hosts in the transmission chain, a partition $\mathfrak{H}$ of $E(\mathcal{T})$ with $l$ parts may assign more than one tip to each host, and a set $\mathbf{I}$ of intervals place limits on the timings of the infections of the sampled hosts. (The unsampled hosts are not given time limits.) Let $\mathbf{D}_{(m,n)}$ be the set of ways in which $m$ unlabelled objects can be placed in $n$ labelled, possibly empty containers. We express a partition $\mathfrak{D}\in\mathbf{D}_{(g,h)}$ as a function $\mathfrak{D}:\{1,\ldots,n\}\to\{0,\ldots,m\}$ taking the index of each container to the number of elements it contains. This is used when summing over the ways that $g$ remaining unsampled parts can be distributed amongst a node's $h$ child subtrees.

The numbers we require are, for a node $u$ of $\mathcal{T}$:

- $|\mathbf{P}_m(\mathcal{T}^*_u;\mathfrak{H},\mathbf{I}\cup[\gamma,\infty))|$, counting the partitions of $\mathcal{T}^*_u$. If $\mathcal{T}_u$ has one tip, which is inside its own element of $\mathbf{I}$, the this is equal to 1 if $m=0$ and 0 otherwise.

- $|\mathbf{PU}_m(\mathcal{T}_u;\mathfrak{H},\mathbf{I})|$, counting the partitions of $\mathcal{T}_u$ where the root is in an unsampled part. If $\mathcal{T}_u$ has one tip then this is always 0.
- For all $i \in \{1,\dots,l\}$, $|\mathbf{P}^i_m(\mathcal{T}_u;\mathfrak{H},\mathbf{I})|$, counting the partitions of $\mathcal{T}_u$ where the root shares a part with the tips in $H_i$. If $\mathcal{T}_u$ has one tip ($u$ itself), which is inside its own element of $\mathbf{I}$, the this is equal to 1 if $m = 0$ and $u \in H_i$ and 0 otherwise.
- $|\mathbf{PS}_m(\mathcal{T}_u;\mathfrak{H},\mathbf{I})|$, which is equal to $\sum_{i=1}^{l} |\mathbf{P}^i_m(\mathcal{T}_u;\mathfrak{H},\mathbf{I})|$
- $|\mathbf{P}_m(\mathcal{T}_u;\mathfrak{H},\mathbf{I})|$, which is equal to $|\mathbf{PU}_m(\mathcal{T}_u;\mathfrak{H},\mathbf{I})| + |\mathbf{PS}_m(\mathcal{T}_u;\mathfrak{H},\mathbf{I})|$

Suppose $u$ has $n$ children $uC_1,\dots,uC_n$. No novel arguments are needed for the following results, which are just the result of careful accounting:

**Proposition 14.**

$$|\mathbf{P}_m(\mathcal{T}^*_u;\mathfrak{H},\mathbf{I}\cup[\gamma,\infty))| = \begin{cases} |\mathbf{P}_m(\mathcal{T}_u;\mathfrak{H},\mathbf{I})|, & h(u) < \gamma \text{ or } u \text{ is a bridge node} \\ |\mathbf{P}_m(\mathcal{T}_u;\mathfrak{H},\mathbf{I})| + \sum_{\mathfrak{D}\in\mathbf{D}_{(m,n)}} \left(\prod_{i=1}^{n} |\mathbf{P}_{\mathfrak{D}(i)}(\mathcal{T}^*_{uC_i};\mathfrak{H},\mathbf{I}\cup[\gamma,\infty))|\right), & \text{otherwise} \end{cases}$$

**Proposition 15.**

$$|\mathbf{PU}_m(\mathcal{T};\mathfrak{H},\mathbf{I})| = \begin{cases} \sum_{\mathfrak{D}\in\mathbf{D}_{(m-1,n)}} \left(\prod_{i=1}^{n} |\mathbf{P}_{\mathfrak{D}(i)}(\mathcal{T}^*_{uC_i};\mathfrak{H},\mathbf{I}\cup(-\infty,\infty))|\right), & u \text{ is not a bridge node} \\ 0, & u \text{ is a bridge node} \end{cases}$$

**Proposition 16.** *Let $B \subseteq \{1,\dots,n\}$ be such that for $j \in B$, $uC_j \in a(H_i)$ and let $B'$ be its complement. If $B$ has more than one element, $u$ is a bridge node. Then:*

$$|\mathbf{P}^i_m(\mathcal{T}_u;\mathfrak{H},\mathbf{I})| = \begin{cases} \sum_{\mathfrak{D}\in\mathbf{D}_{(m,n)}} \left(\prod_{j\in B} |\mathbf{P}^i_{\mathfrak{D}(j)}(\mathcal{T}_{uC_j};\mathfrak{H},\mathbf{I})| \times \prod_{k\in B'} |\mathbf{P}_{\mathfrak{D}(k)}(\mathcal{T}^*_{uC_k};\mathfrak{H},\mathbf{I}\cup[\alpha_i,\infty))|\right), & h(u) \in I_i \text{ and } u \notin \mathbf{b}(H_p) \text{ for } p \neq i \\ 0, & \text{otherwise} \end{cases}$$

The sampling traversal is analogous to that described under "Incomplete sampling" above, and the random assignment of unsampled hosts with no corresponding parts of the partition can also be done exactly as described in that section.
