## Supplementary material for "Transmission trees on a known pathogen phylogeny: enumeration and sampling"

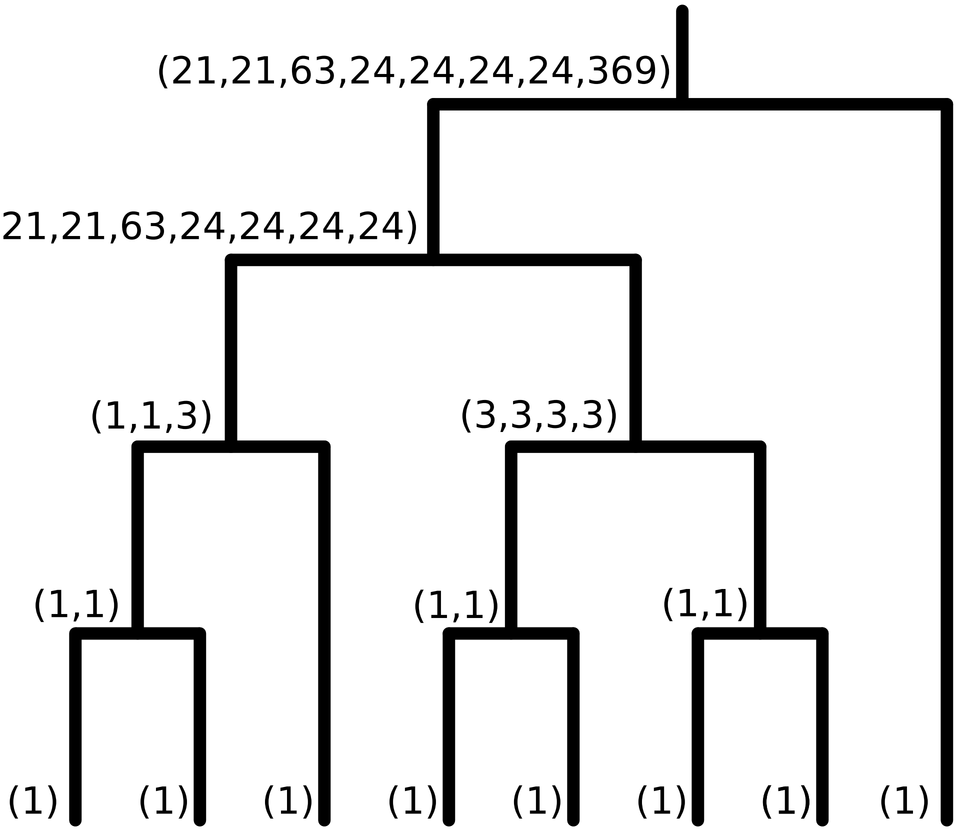


Figure S1: The calculation of all $|\mathbf{P}^i(\mathcal{T})|$ if $\mathcal{T}$ is the tree in figure 4. Each internal node $u$ rooting a subtree $\mathcal{T}_u$ is annotated with a tuple of the nonzero values of $|\mathbf{P}^i(\mathcal{T}_u)|$, appearing in the same order as the subtree's tips as displayed from left to right.

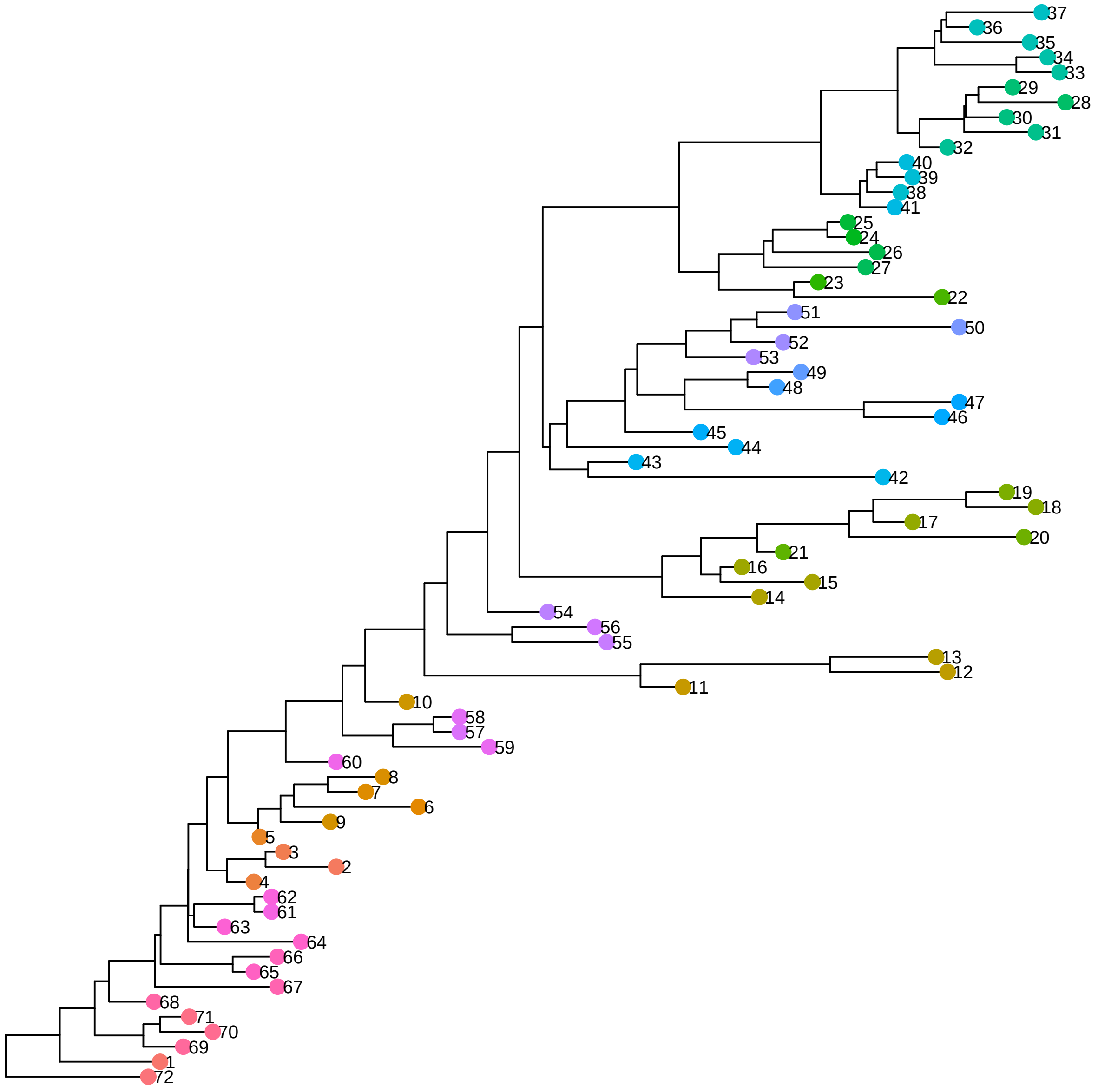


Figure S2: The 72-tip phylogeny of TB isolates used to compare *TransPhylo* and *STraTUS*. Colours and numbers are the same as those in figure 10.

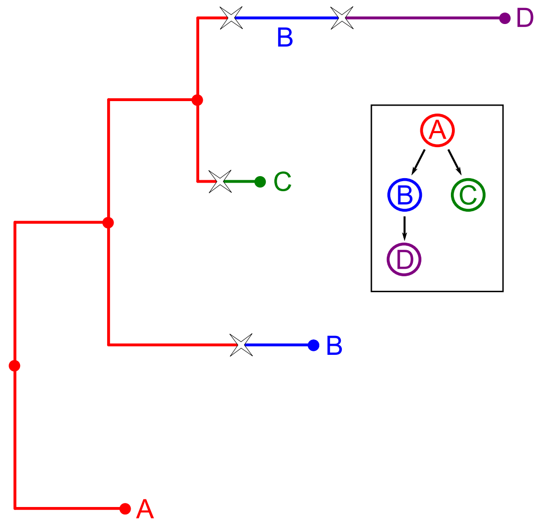


Figure S3: For this phylogeny, the inset transmission tree is impossible under the complete bottleneck assumption, but possible if it is relaxed and host A can transmit two lineages to host B, as shown. Stars represent transmission events.

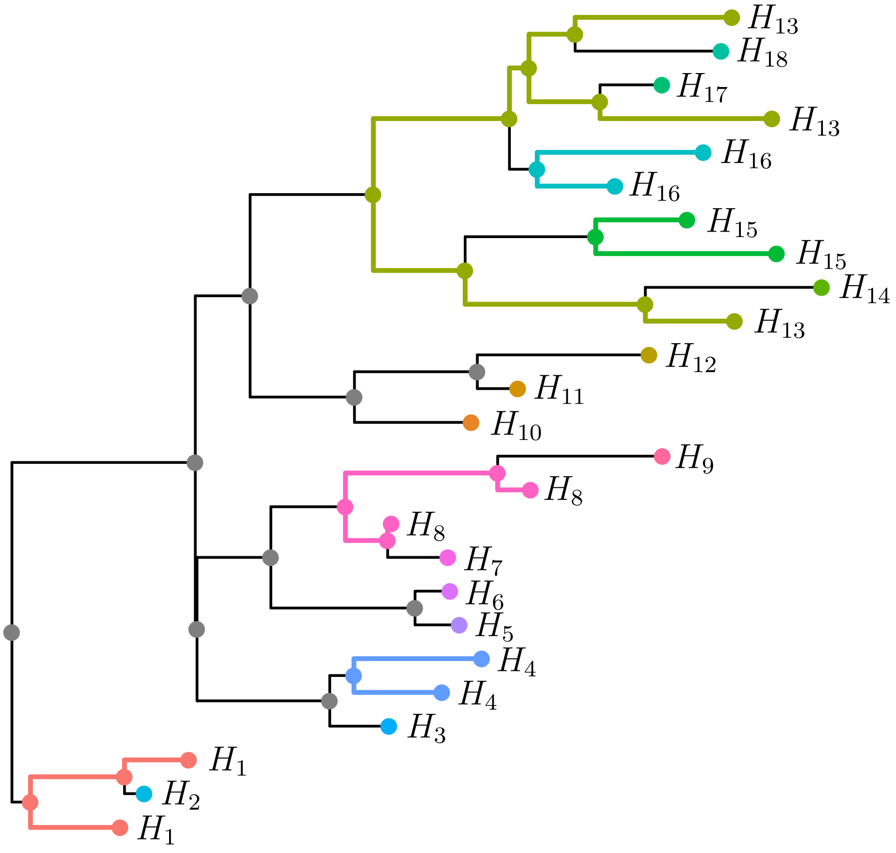


Figure S4: An illustration of bridges. The partition $\mathfrak{H}$ of the tips of this tree has eighteen parts $\{H_1, \ldots, H_{18}\}$, and each tip is labelled, and coloured, according to the part it belongs to. Six parts ($H_1$, $H_4$, $H_8$, $H_{13}$, $H_{15}$ and $H_{16}$) contain more than one tip, and define bridges that contain internal nodes. For each such $H_i$ the nodes in the bridge $\mathbf{b}(H_i)$, and the branches connecting them, appear as the same colour as the elements of $H_i$. Nodes that are not members of any bridge are grey.
